# Structural Dynamics of IRE1 and its Interaction with Unfolded Peptides

**DOI:** 10.1101/2025.02.28.640832

**Authors:** Elena Spinetti, Grzegorz Ścibisz, G. Elif Karagöz, Roberto Covino

## Abstract

The unfolded protein response (UPR) is a crucial signaling network that preserves endoplasmic reticulum (ER) homeostasis, impacting both health and disease. When ER stress occurs, often due to an accumulation of unfolded proteins in the ER lumen, the UPR initiates a broad cellular program to counteract cytotoxic effects. Inositol-requiring enzyme 1 (IRE1), a conserved ER-bound protein, is a key sensor of ER stress and activator of the UPR. While biochemical studies confirm IRE1’s role in recognizing unfolded polypeptides, high-resolution structures showing direct interactions remain elusive. Consequently, the precise structural mechanism by which IRE1 senses unfolded proteins is debated. In this study, we employed advanced molecular modeling and 136,7 μs of atomistic molecular dynamics simulations to clarify how IRE1 detects unfolded proteins. Our results demonstrate that IRE1’s luminal domain directly interacts with unfolded peptides and reveal how these interactions can stabilize higher-order oligomers. We provide a detailed molecular characterization of unfolded peptide binding, identifying two distinct binding pockets at the dimer’s center, separate from its central groove. Furthermore, we present high-resolution structures illustrating how BiP associates with IRE1’s oligomerization interface, thus preventing the formation of larger complexes. Our structural model reconciles seemingly contradictory experimental findings, offering a unified perspective on the diverse sensing models proposed. We elucidate the structural dynamics of unfolded protein sensing by IRE1, providing key insights into the initial activation of the UPR.

## Introduction

The endoplasmic reticulum (ER) is central for protein folding, maturation, and delivery of membrane and secreted proteins in eukaryotic cells.^1^ The lumen of the ER presents unique features, such as high calcium ion concentration,^2^ oxidizing redox potential, ^3^ and a high number of chaperones.^4^ These unique features must be preserved to ensure correct protein production. Loss of ER homeostasis can lead to the accumulation of unfolded and misfolded proteins, a phenomenon known as ER stress.

The Unfolded Protein Response (UPR)^5^ is a collection of signaling pathways that allows eukaryotic cells to counteract ER stress. The membrane protein inositol-requiring enzyme 1 (IRE1) is the UPR’s central transducer in all eukaryotes and the most evolutionarily conserved branch of the UPR.^6,7^ It activates its signaling pathway by forming oligomers and large supra-molecular assemblies.^8–11^ IRE1 is a single-pass transmembrane protein and can sense the unfolded protein load of the ER lumen through its core luminal domain (cLD).^12–14^ Accumulation of unfolded proteins in the ER promotes the formation of multimeric assemblies, allowing the cytosolic kinase domains to trans-phosphorylate^15^ and activate the RNase domains. This, in turn, initiates splicing of *XBP1* mRNA, which encodes for a transcription factor,^16,17^ leading to the transcription of UPR target genes. In addition, the active RNase initiates mRNA decay.^18,19^ The UPR aims at restoring ER homeostasis, and if stress is not mitigated in a timely manner, the UPR triggers cell death via apoptosis. ^20^ Chronic ER stress has been implicated in various diseases, from diabetes to cancer.^21^ Thus, maintaining the folding equilibrium in the ER is crucial for the proper functioning and viability of eukaryotic cells.

The cLD initiates IRE1’s activation, yet the mechanism by which it detects the accumulation of misfolded proteins remains debated. Two main models have been proposed, which differ in whether the cLD senses unfolded proteins directly or indirectly. ^22^ In the direct binding model, IRE1’s cLD interacts directly with unfolded proteins, triggering assembly and activation.^23^ Observations that the crystal structure of the cLD dimer^17^ of *Saccharomyces cerevisiae* (yIre1) presents a groove in the central area that resembles the major histocompatibility complex class I (MHC-I) peptide-binding cleft^12^ have led to the hypothesis that IRE1 uses a mechanism for unfolded peptide binding similar to the antigen-presenting protein. Subsequently, the binding of unfolded peptides to the yIre1 cLD was demonstrated in vitro and in cells.^13^ According to this hypothesis, the peptides bind to the groove delimited by the two monomers’ helices at the center of the dimer, spanning the dimeric interface. However, the structure of human IRE1 ubiquitous isoform *α* (hIRE1*α*) cLD dimer features a narrower cleft than the yeast Ire1 cLD.^24^ These observations inspired the hypothesis that the conformation observed in yeast represents an ‘open’ state of the dimer, whereas the crystal structure of human cLD represents a ‘closed’ dimer state that requires opening to accommodate peptides.^14^ Recently, evidence of direct binding of unfolded polypeptides to the human IRE1*α* cLD central groove accumulated,^14,25–28^ indicating that this domain preferably binds peptides enriched in aromatics, hydrophobics, and arginine residues. Moreover, new experiments showed that the human IRE1 already forms dimers in non-stress conditions,^29^ suggesting that peptides might bind preexisting dimers and induce oligomerization required for the subsequent activation of the sensor. Despite this evidence, we still lack a clear structural understanding of the direct sensing model and how the binding of unfolded peptides promotes the formation of higher-order assemblies.

In the indirect activation model, BiP, a chaperone of the Hsp70 family in the ER lumen, mediates interactions with unfolded proteins.^30,31^ In this scenario, BiP binds to IRE1, preventing it from forming dimers and oligomers in the absence of stress. At the onset of ER stress, BiP binds to accumulating unfolded proteins, leaving IRE1 free to self-associate and start the UPR signaling cascade. BiP could bind to IRE1 either via its ATPase domain or substrate-binding domain (SBD).^32^ Interaction through the substrate domain would hint at a direct competition between IRE1 and unfolded proteins, while binding through the nuclease domain supports an allosteric mechanism.^32^ Moreover, BiP binds to regions of IRE1, which are disordered and part of the oligomerization interface.^25,26^

In recent years, the field has been converging towards integrating the direct and indirect binding models, where unfolded proteins and BiP interaction with cLD both regulate IRE1.^5^ We aimed to integrate evidence about the human IRE1*α* cLD early activation phase to develop a structural understanding of this protein domain’s sensing mechanism.

Here, we performed multi-microsecond atomistic molecular dynamics (MD) simulations to investigate the stability of the human IRE1*α* cLD dimer, its structural features, and its interaction with unfolded polypeptides, and with BiP. By integrating the crystal structure with Alphafold Multimer^33,34^ prediction, we identified the *β*-sheet interface as well as hydrophobic side chains orientation as the key elements of a stable dimer. Comparison of the dynamics of yeast and human cLD dimer highlighted differences, suggesting alternative stress-sensing strategies. Our simulations provided structural evidence that unfolded polypeptides bind directly to the center of the human IRE1*α* cLD dimer, reinforcing their crucial role in cluster formation during the initial phases of ER stress. Additionally, we provide structural evidence supporting a model in which hIRE1*α* cLD is bound to BiP at the oligomerization interface, inhibiting cluster formation while allowing dimer formation.

## Results

### The hIRE1 *α* cLD forms a stable dimer

An accurate determination of proteins’ structural dynamics is crucial to understanding their function. MD simulations use the laws of physics to simulate the movements of atoms in a protein, generating a trajectory, a “movie” showing how its structure changes in time. The trajectory can reveal conformational changes, alternative structural organizations, and their relative stability. MD simulations of multiple molecules can reveal how they interact and whether they form stable assemblies. MD simulations require an initial structural model to start generating a trajectory. The human IRE1*α* core luminal domain (cLD) structure has been resolved by X-ray crystallography and deposited in the Protein Data Bank as a symmetric homodimer (PDB ID 2HZ6^24^). This structure lacks residues in two disordered regions: DR1 (residues L131-S152) and DR2 (residues P308-N357). We used protein sequence information and modeled these gaps as unstructured loops, obtaining the “PDB model” (**Fig. 1A**, modeled regions are DR1 in cyan and DR2 in orange). Despite its relatively high resolution of 3.10 Å, the PDB structure shows low percentile scores in the wwPDB (Worldwide Protein Data Bank) ^35^ validation report for global metrics, including torsion angle outliers (Ramachandran and sidechain) and quality of fit (RSRZ outliers). Notably, the PDB dimeric assembly was derived from the crystal packing of the asymmetric unit of the crystal, which includes only one monomer.

**Figure 1.**
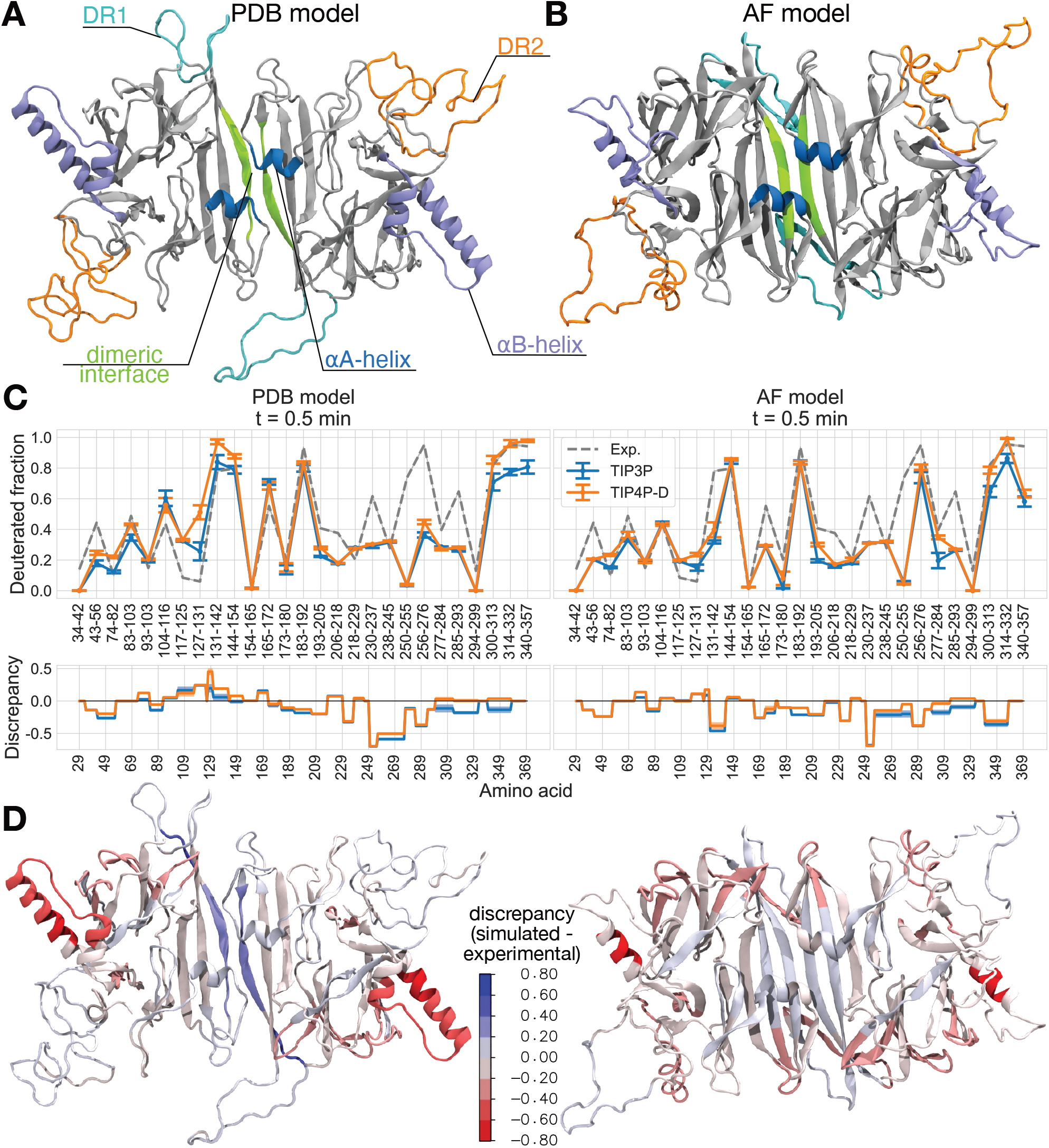
The hIRE1*α* cLD forms a stable dimer validated by hydrogen-deuterium exchange experimental data. (A) The PDB model: structural model of hIRE1*α* cLD dimer as obtained by Zhou et al.^24^ (PDB ID 2HZ6) with the addition of missing residues, in particular DR1 (cyan) and DR2 (orange). Backbone in cartoon representation, areas of interest highlighted in different colors. (B) The AF model: AlphaFold 2 Multimer prediction of hIRE1*α* cLD dimer (P29-P368). Representation style as in Fig. 1A.(C) The deuterated fraction obtained from experimental results (dashed line, shaded area indicates the error we calculated from bootstrapping) published by Amin-Wetzel et al.^25^ and the fraction computed from MD simulations (solid lines, blue for TIP3P water and orange for TIP4PD water) for the PDB and AF model at incubation time point 0.5 min. This time point corresponds to experimental incubation times, not MD simulation time. Each point represents the mean value derived from three replicas and two monomers per replica. The error bars were obtained from bootstrapping. Below each absolute value plot, we report the discrepancy, which is defined as the difference between the simulated and experimental deuterated fractions, with the shaded area indicating the corresponding error. (D) Visualization of the discrepancy values from Fig. 1C onto the molecular structures of the PDB and AF model simulated in TIP4P-D water (TIP3P shown in Fig. 10A). Shades of blue indicate regions where the simulated structure is more flexible and solvent-accessible than observed experimentally, while shades of red indicate regions where the simulated structure is more rigid and less solvent-accessible than expected.

We generated an alternative cLD dimer structure using AlphaFold 2 Multimer,^33,36^ termed the “AF model” (**Fig. 1B**). The PDB and AF models were similar, with an RMSD of 3.34 Å (aligned on the *β*-sheet floor, excluding DR1 and DR2) **(Supplementary Fig. 7A**). Notable structural features (**Fig. 1A-B**) include the dimer interface (green), where the two monomers establish contacts through an antiparallel *β*-sheet interaction that determines the cyclic C2 symmetry of the dimer; the MHC-like groove delimited by *α*A-helices (blue), and the eight-stranded *β*-sheet floor. AlphaFold captured these features while revealing slight structural differences: DR1 was partly predicted as a *β*-hairpin rather than disordered (cyan), and DR2 (orange) featured an *α*-helix (K349-L361). Additionally, the *α*B-helices (violet) were shorter in the AF model than in the PDB model. The discrepancies in these regions were intriguing, as the disordered regions are difficult to study experimentally and might contain transient secondary structures. At the same time, the *α*B-helices are involved in the formation of a dimer-dimer interface.^14^ A comparison between the AlphaFold 2 and AlphaFold 3 predictions provided a very similar hIRE1*α* cLD dimer model (**Supplementary Fig. 7B**). All five AlphaFold 2 predictions closely resembled the top-ranked model used for our simulations (**Supplementary Fig. 7C**). In contrast, the five AlphaFold 3 predictions yielded greater variability in DR2 organization and longer helices in DR2, but still consistently maintain low pLDDT scores in this region, indicating disorder (**Supplementary Fig. 7D**).

Current all-atom force fields used in MD simulations are mainly designed to reproduce the dynamics of folded and globular proteins. ^37^ They are less accurate in simulating the structural ensemble of disordered proteins and regions, usually describing structural ensembles that are excessively compact.^38,39^ One culprit is the commonly used water model called TIP3P,^40^ which tends to underestimate London dispersion interactions significantly.^38^ The TIP4P-D water model was developed to address limitations of existing force fields in reproducing the structural ensembles of intrinsically disordered proteins and regions. It incorporates enhanced dispersion and moderately stronger electrostatic interactions to improve the balance between water dispersion and electrostatics.^38^ Zapletal et al.^39^ showed that for proteins containing both folded and disordered regions, the CHARMM36m force field^41^ in combination with the TIP4P-D water model provides a robust framework, preventing collapse of disordered regions while preserving folded regions. Acknowledging that the behavior of disordered regions can be case-specific, we conducted MD simulations of the two cLD dimer models using the CHARMM36m force field with both TIP3P and TIP4P-D water models.

During our 2 μs-long simulation replicas, the AF and PDB models of the dimer always remained stable (**Supplementary Fig 7E-F**), even if the PDB model presented some instabilities in one of the central hydrogen bonds (**Supplementary Fig 8**). An initial PDB model with modified side chain orientations in residues L116 and Y166 due to the modeling of neighboring missing DR1, caused the dimer to dissociate in one-third of the replicas (**Supplementary Fig. 9A-B**). The proximity of these residues to W125 and P108, essential for dimer formation,^24,25^ highlights the importance of hydrophobic residues in stabilizing the dimer interface. The final PDB model, with correctly oriented L116 and Y166 (**Supplementary Fig. 9B**), was stable in simulations in both TIP3P and TIP4P-D water (**Supplementary Fig. 7E**). While modeling missing regions carries inherent risks, it highlighted the critical roles of L116 and Y166 and the value of testing alternative models. The AF model showed high stability of the dimer interface, and the DR1 fold might contribute to this stabilization. Consequently, we modified the AF model by substituting DR1 with an unfolded loop (**Supplementary Fig. 9C**). We observed no differences in the stability of the dimer interface, concluding that the DR1 *β*-hairpin fold does not contribute to dimer stability. In summary, we modeled the human IRE1*α* cLD dimer by complementing the crystal structure with disordered regions and utilizing structure prediction for an alternative model.

Our simulations showed that the dimers remained consistently stable, agreeing with recent experimental results suggesting the IRE1*α* might constitutively be in a dimer organization even in the absence of stress.^29^

### Hydrogen-deuterium exchange experimental data validate the cLD dimer structure

The two models of human IRE1*α* cLD dimer exhibited structural differences in certain regions, showing variations in secondary structures and flexibility. Furthermore, the two distinct water models used, TIP3P and TIP4PD, led to differences in flexibility observed for the same structural model. Therefore, we aimed to validate our models by comparing them with experimental data. In the study by Amin-Wetzel et al., ^25^ the authors published the results of HDX-MS (hydrogen(^1^H)-deuterium(^2^H) exchange mass spectrometry) experiments on the human IRE1*α* luminal domain. In HDX-MS experiments, a protein is incubated in a deuterated buffer, permitting the amide ^1^H from the protein backbone to exchange with ^2^H from the solution.^42^ The exchange rate is influenced by the exposure of the backbone hydrogen to the aqueous environment, determined by the amino acid sequence’s chemical nature, the protein’s folded state, and its dynamic rearrangements and flexibility. A hydrogen atom in a folded region of the protein will be less likely to exchange than one in an unstructured region. After incubating the protein in solution for a designated period of time, it is enzymatically digested, and the deuterium uptake of each fragment is assessed through mass spectrometry. These experiments provide insights into the flexibility and solvent accessibility of the protein regions. Specifically, the measured deuterated fraction of a peptide correlates with the flexibility of the corresponding protein region and is inversely related to its folded structure. We calculated the theoretical deuterated fraction from our simulations using the method by Bradshaw et al.^43^ and compared it to the experimental data (**Fig. 1C-D and Supplementary Fig. 10**) (see Methods). By juxtaposing the experimental data with the results from our PDB and AF models, we can confirm the accuracy of each model.

Both structural dimer models agree well with the HDX-MS data (**Fig. 1C**), even though only a mixture of the two models could recapitulate the experimental data entirely (**Fig. 1D and Supplementary Fig. 10-11**). The prediction of the deuterated fraction for DR1 based on the PDB model aligns most closely with experimental data (**Supplementary Fig. 11**). This suggests that DR1 is disordered, contrary to the folding predictions made by the AF model. The dimeric interface was most accurately represented by the AF model, suggesting that the stable dimeric conformation could be constitutive under experimental conditions. The dimeric interface of the PDB model had a predicted deuterated fraction higher than the experimental one, indicating a region that is more water-accessible and flexible than expected. This corresponds to the instability of one H-bond observed in simulations and mentioned earlier (**Supplementary Fig. 8**), likely due to a suboptimal interface in the dimer of Zhou et al.^24^ On the other hand, the AF model has a better-formed interface due to optimization by AlphaFold, and the hydrogen bonds were stable throughout all simulations. The *α*B-helices (V245-I263) are long *α*-helices found in the crystal structure of human IRE1*α* cLD that are not found in the yeast homolog structure.^12,24^ This helical structure may inhibit oligomer formation, as suggested by Zhou et al., ^24^ and a structural rearrangement might be necessary for oligomerization, as supported by Nuclear Magnetic Resonance (NMR) spectroscopy experiments. ^14^ In the PDB model, these long helices showed a predicted deuterated fraction lower than the experimental value. This result indicates that the region is slightly more structured and less flexible during simulations than expected. This may be because contacts within the crystal stabilize longer *α*B-helices compared to what is found in solution. Conversely, the *α*B-helices are more unfolded in the AF model, and our analysis suggested that this structure aligned better with the experimental data. This supports the idea that the *α*B-helices could be shorter than what was found in the crystal, and this would allow the structural arrangements observed by the NMR experiments.^14^

Overall, different protein regions are best represented by different protein models (**Supplementary Fig. 11**). However, the AF model has the most stable dimeric interface, and in TIP4P-D water, it produces the highest number of deuterated fractions closer to the experimental ones (**Supplementary Fig. 11A**). Therefore, we chose to use the AF model of the dimer for all our subsequent simulations.

### The putative groove of human IRE*α* cLD is dynamic but unable to contain peptides

The fact that the Ire1 cLD dimer closely resembles the peptide-binding groove in the MHC-I, in which peptides nestle into a central cleft, suggested a model for interactions of unfolded proteins and Ire1 in *Saccharomyces cerevisiae*. However, Zhou et al.^24^ argued that the human conformation of the dimer has a groove that is too narrow to accommodate peptides. Indeed, the groove width varies significantly between the two structures in the Protein Data Bank: the groove width is 6.8 Å in hIRE1*α* and 10.7 Å in yIre1 (yIre1 cLD: PDB ID 2BE1^12^). (**Fig. 2A-C and Supplementary Fig. 12A**). In our simulations, the PDB model of the hIRE1*α* cLD dimer exhibited a less stable dimeric interface. Consequently, we compared the yeast dimer with the AF model of the human dimer, which features a groove width similar to that of the PDB model, measuring 7.4 Å. In simulations of the dimeric structures, the average groove width was 7.3 ± 0.1 Å for the human cLD and 8.9 ± 0.1 Å for the yeast cLD, averaged over three TIP3P and three TIP4P-D replicas per system (**Fig. 2C**). In both humans and yeast, we noted a fluctuating groove width, which was somewhat wider in yeast. Nonetheless, we were unable to differentiate between the open and closed conformations in either structure. These data suggest, in contrast to what has been proposed before, that human IRE1*α* cLD in isolation does not adopt an open conformation.^14,44^

**Figure 2.**
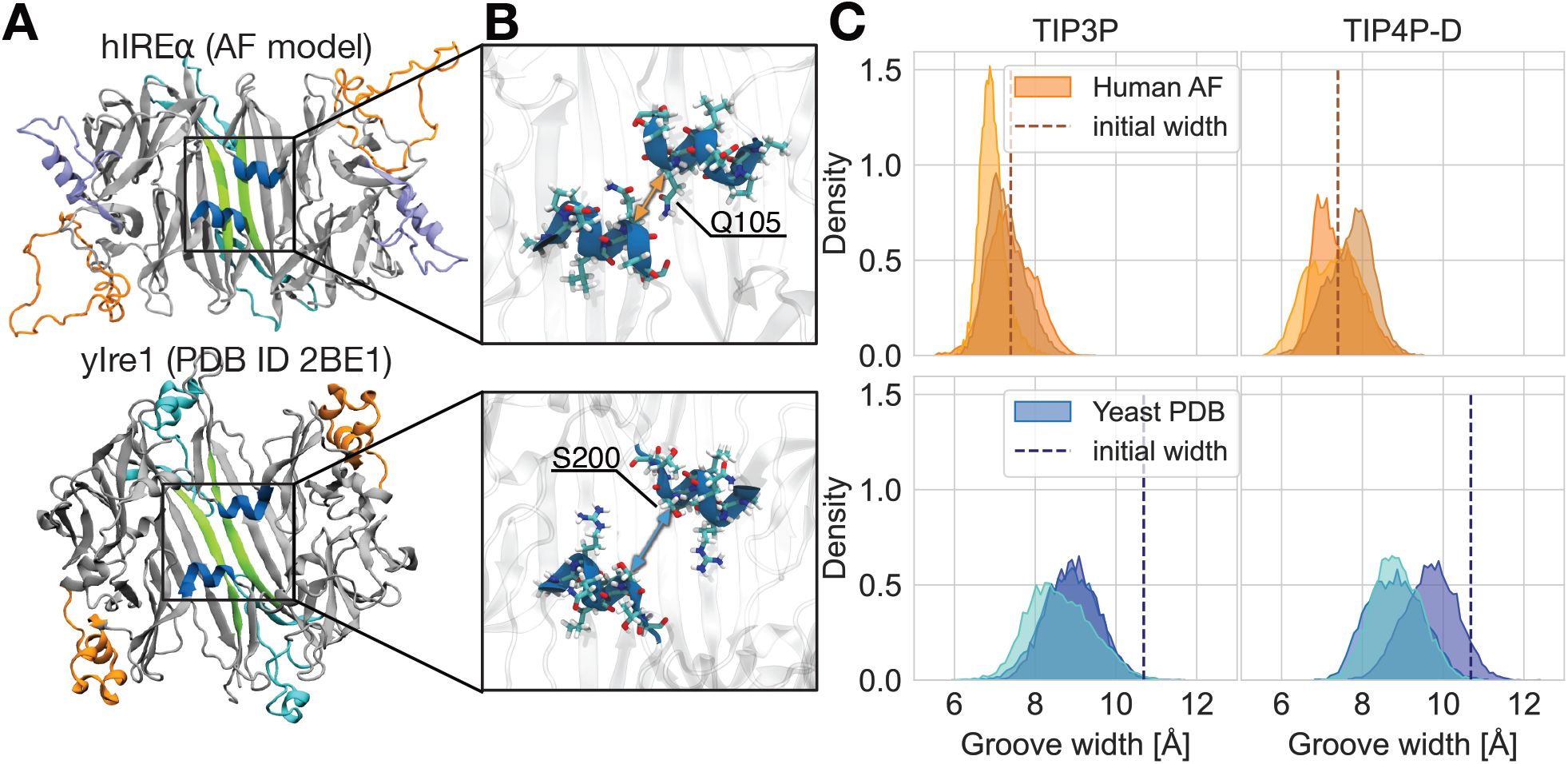
The human IRE1*α* cLD dimer has a narrower and more occluded central groove than its yeast homolog. (A) Top panel: structure of human IRE1*α* (hIRE1*α*) cLD dimer (AlphaFold model). Bottom panel: structure of the S. Cerevisiae IRE1 (yIre1) cLD dimer (PDB ID 2BE1). (B) Close-up on hIRE1*α* (top panel) and yIre1 (bottom panel) structures on the central groove delimited by the *α*A-helices. Helices delimiting the groove are depicted in blue cartoon and side chains in stick representation, while the rest of the dimer is transparent. The arrows indicate the distance between C*α* atoms used to compute the groove width. (C) Probability density distribution of the groove width of: hIRE1*α* cLD dimer computed between C*α* of Q105 during simulations in TIP3P and TIP4P-D water (top panel); yIre1 luminal domain dimer computed between C*α* of S200 during simulations in TIP3P and TIP4P-D water (bottom panel). The different shades of colors in the distributions represent the three system replicas for each water model. Dashed lines indicate the groove width of the dimer model prior to energy minimization.

Despite the structural similarities in the central regions of human and yeast cLD (**Fig. 2A**), a closer examination reveals differences in the side chains of the *α*A-helices. In yeast, these are serines (S200), while in humans, they are glutamines (Q105) (**Fig. 2B**). The glutamines in human cLD possess a longer side chain than serines. The configurations they adopt during simulations enable them to occupy the space between the *α*A-helices (**Supplementary Fig. 12B**). These side chain arrangements result in spatial occupancy of the groove, particularly in instances where the groove width is narrow, as observed here.

In conclusion, our simulations revealed that in contrast to the yeast cLD that features a wider groove, the putative central groove of human IRE1*α* cLD is predominantly narrow and occluded by the steric obstruction from neighboring side chains. As our MD simulations did not show large conformational changes leading to the opening of the groove, our data suggest that it is unlikely that peptides bind into the central part of the MHC-like groove in the human IRE1*α* cLD.

### Unfolded polypeptides bind to hIRE1*α* cLD dimer surface

Despite its structural diversity compared to MHC-like cleft, hIRE1*α* cLD can directly bind to polypeptides *in vitro*.^14,26^ Therefore, we aimed to investigate the characteristics of peptides in complex with the hIRE1*α* cLD dimer model through MD simulations. Our goal was to elucidate a potential binding pose and identify the relevant features of unfolded proteins and the cLD that affect the binding. The simulated complexes were initialized by positioning the peptides in a fully stretched conformation above the central region of the dimer (**Fig. 3A-C, at t = 0 μs**). This enabled the peptides to equilibrate near the surface of hIRE1*α* within a few nanoseconds. In initial simulations with peptides valine8 and MPZ1-N, we positioned the polypeptides over the cLD, aligning them parallel to the principal axis of the central groove in accordance with the proposed binding mode. We refer to this pose as the “0° orientation”, as the peptide forms a 0 ° angle with the principal axis of the groove. We observed that the peptides could rearrange into an orientation perpendicular to the central groove axis, while maintaining contact with the dimer (**Fig. 3A, Supplementary Fig.13A**, valine8 TIP4P-D, and **Supplementary Fig.14**). Conversely, when MPZ1-N was initially oriented perpendicularly to the groove, it did not transition to a parallel (0°) orientation (**Supplementary Fig.14**). We refer to these poses as the “90° orientation” and “270° orientation”. This finding suggests that the perpendicular binding mode is preferred. Therefore, we adopted this orientation as the baseline for subsequent simulations.

**Figure 3.**
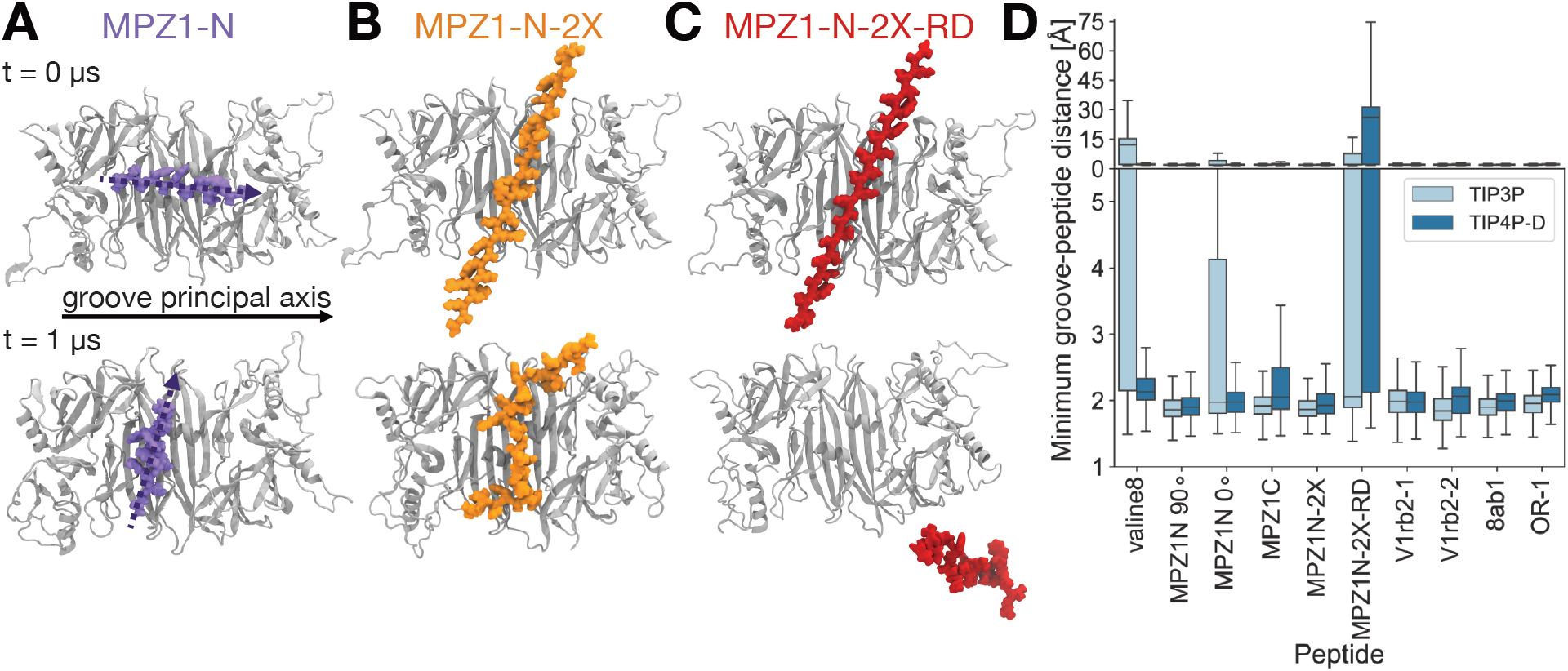
Unfolded peptides can bind with specificity to the center of the cLD dimer, even if the groove is closed. (A) AF model of the cLD dimer (gray cartoon) simulated with the unfolded peptide MPZ1-N (violet) positioned parallel to the principal axis of the central groove (t = 0 μs). During the simulation, MPZ1-N rearranges to a vertical position (t = 1 μs). The direction of the principal axis of the central groove is represented by a solid arrow, while the direction of the principal axis of the peptide is indicated by a dashed arrow. (B) AF model of the cLD dimer simulated with unfolded peptide MPZ1N-2X (orange), which stably binds to the center of the dimer. (C) AF model of the cLD dimer simulated with unfolded peptide MPZ1N-2X-RD (red), which unbinds in half of the replicas. (D) Box plot distributions of the minimum groove-peptide distance. Each box plot reports on the results from 3 replicas (in TIP3P or in TIP4PD) of a system containing a dimer and one of the unfolded peptides.

We tested the stability of cLD complexes with polypeptides that were previously characterized by fluorescence anisotropy experiments^14,26,45^ (**Fig. 3B-D and Supplementary Fig. 13A-B**). These peptides were primarily derived from Myelin Protein Zero (MPZ) and displayed a range of binding affinities (*K*_*1/2*_; **Supplementary Tab. 2**). The most potent binders, MPZ1-N, MPZ1N-2X, and 8ab1, remained bound to the dimer throughout all the 1 μs-long simulation replicas in TIP3P and TIP4P-D water. In contrast, we observed partial dissociation of MPZ1-C, which is a weaker binder. All PERK-targeted peptides, namely OR-1, V1rb2-1, and V1rb2-2, demonstrated strong binding to the cLD dimer. The apolar peptide valine8 dissociated in one-third of the simulations. By mutating all the arginines in MPZ1N-2X to aspartic acid, we obtained the MPZ1N-2X-RD. The MPZ1N-2X-RD has six negative charges instead of the six positive charges present on MPZ1N-2X. These mutations led to the dissociation of MPZ1N-2X-RD in half of the simulations, recovering the impaired binding observed in experiments.

Notably, none of the peptides caused meaningful structural changes in the dimer, and we did not observe a groove opening induced by the peptides (**Supplementary Fig. 15A**). However, the unfolded peptides, particularly those rich in positively charged residues, remained bound to the hIRE1*α* cLD dimer surface (**Supplementary Fig. 15B**) and were preferentially oriented perpendicular to the groove’s main axis. AlphaFold3 predictions of the complexes indicate that the peptides adopt the same preferred orientation, despite being predominantly helical (**Supplementary Fig. 16A**). We further assessed the MPZ-derived peptide complexes using MM/PBSA free energy calculations over the final 250 ns of each simulation replica (see Methods), finding binding enthalpies consistent with our observations (**Supplementary Fig. 16B**). In particular, MPZ1N-2X exhibited the lowest binding energy, whereas MPZ1N-2X-RD showed the highest. The simulations generally aligned with the experimental results, establishing a reliable framework for analyzing key binding sites.

### Point mutations destabilize unfolded peptide binding to cLD

The results from polypeptides-cLD simulations indicated that the cLD of hIRE1*α* formed selective interactions with the peptides. To investigate the characteristics of this association, we analyzed the contacts established during the simulations. The peptides interacted with various regions on the surface of the cLD dimer, particularly around the central groove (**Fig. 4A** and **Supplementary Fig. 17A**). Notably, the region above the *α*A-helices played a prominent role in these interactions, primarily due to the presence of a negatively charged residue E102, which might influence selectivity for positively charged peptides. However, the contact analysis primarily reflects the spatial proximity of residues and does not provide information regarding chemical specificity or established bonds. Consequently, these findings offer preliminary insights that necessitate further validation through individual trajectory examinations. Detailed observations of the simulation trajectories indicated that peptides with longer side chains, like arginine, penetrated more into the central region of the dimer. These residues engaged in particular with Y161, D181, and D79, thus forming what we refer to as a polar-charged pocket (**Fig. 4B**, highlighted in yellow). In contrast, a hydrophobic-aromatic pocket defined by Y117, L103, and F98 (**Fig. 4B**, highlighted in red) showed affinity for residues like tryptophan, tyrosine, and leucine. The tyrosines Y117 and Y161 were centrally positioned within the two pockets.

**Figure 4.**
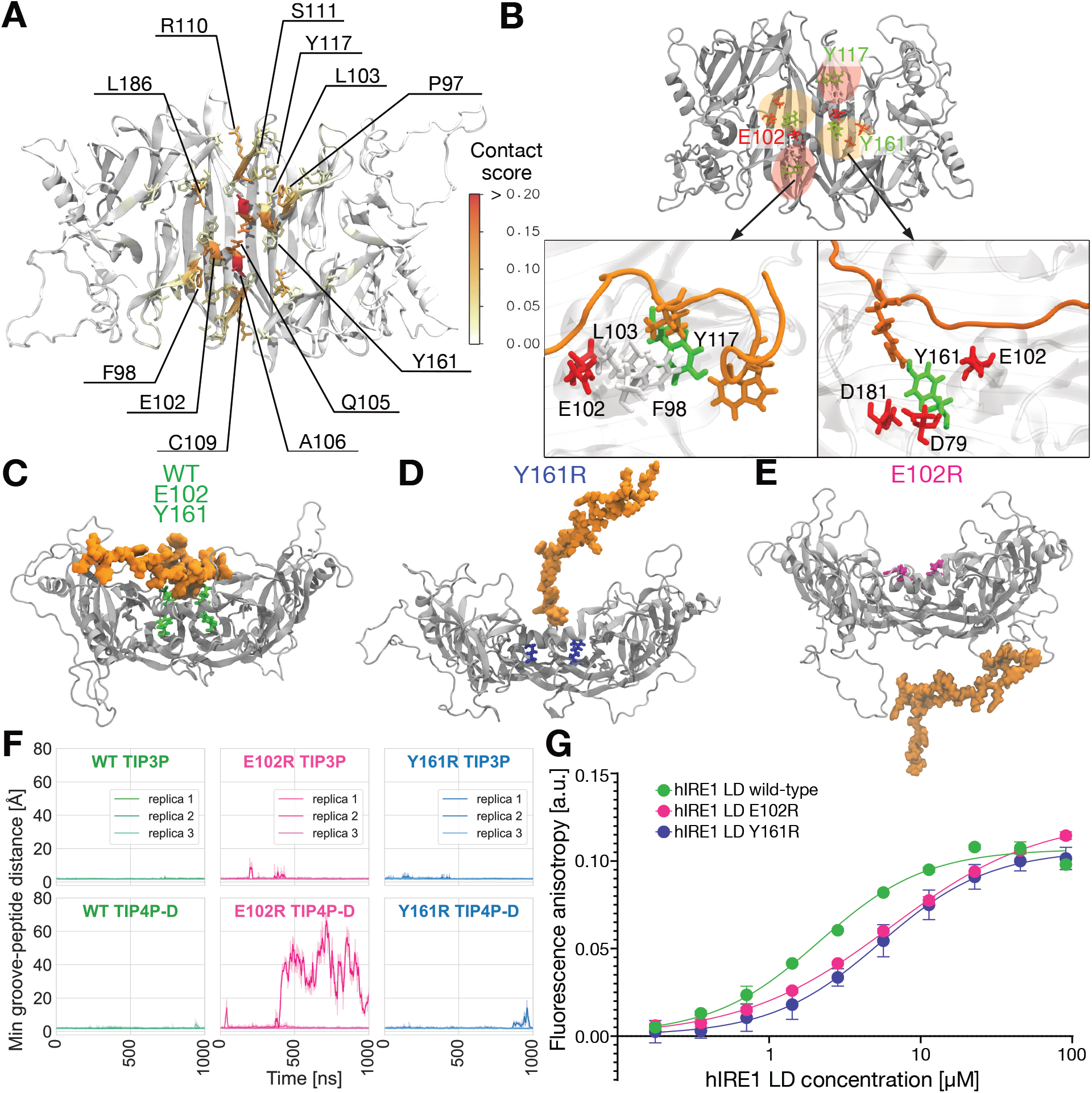
Single point mutations of cLD weaken the binding of unfolded peptide MPZ1N-2X. (A) Analysis of contacts between unfolded peptide and cLD dimer for all MD simulations performed in TIP4P-D water. The contact score reported on the AlphaFold model of the cLD dimer was obtained by computing the contacts from all the simulations, summing them, and normalizing them by the number of frames. Lower values indicate fewer contacts observed. (B) Important areas for peptide binding on the surface of the cLD dimer: the red area identifies a hydrophobic-aromatic binding pocket characterized by residues Y117, F98, and L103; the yellow area identifies a deeper polar-charged binding pocket and is characterized by residues Y161, D181, and D79. Residue E102 on the *α*Ahelices is a hub for contacts on the surface of the dimer. (C) Side view snapshot after 1 μs of simulation of WT hIRE1*α* cLD dimer (gray) in complex with MPZ1N-2X (orange). The amino acids E102 and Y161 on both monomers are represented in green sticks. (D) Side view snapshot after 1 μs of simulation of Y161R hIRE1*α* cLD dimer (gray) in complex with MPZ1N-2X (orange). The amino acid R161 on both monomers is represented in blue stick. (E) Side view snapshot after 1 μs of simulation of E102R hIRE1*α* cLD dimer (gray) in complex with MPZ1N-2X (orange). The amino acid R102 on both monomers is represented in magenta sticks. (F) Time series of the minimum groove-peptide distance for MPZ1N-2X simulated in complex with wild-type, E102R, and Y161R hIRE1*α* cLD dimer in TIP3P (3 replicas) and TIP4P-D (3 replicas) water. The darker lines show the rolling average over 25 frames, while the shaded lines represent the raw data. (G) Fluorescence anisotropy measurements of labeled MPZ1N-2X binding to hIRE1*α* LD wild type and mutants E102R and Y161R.

Thus, we investigated how the point mutations of two key residues, E102R and Y161R, would affect peptide binding by simulating the cLD mutant in complex with MPZ1N-2X (**Fig. 4C-E**). We initialized the systems in the pose described for the other peptide-cLD systems described earlier (**Fig. 3B, t = 0 μs**). In simulations of the wild-type (WT) cLD dimer, the peptide generally remained near the center (**Fig. 4C,F**). By contrast, MPZ1N-2X displayed reduced binding to E102R, fully dissociating in one TIP4P-D replica (**Fig. 4E**,**F**). A similar trend was observed for Y161R, where one partial dissociation event occurred (**Fig. 4D,F**). Comparative analysis of MPZ1N-2X contact sites on the WT and mutant cLD dimers (**Supplementary Fig. 17B-D**) revealed that, in the presence of mutations, the peptide engages a broader surface region rather than remaining centrally localized, while forming fewer contacts with the specific residues (**Supplementary Fig. 18A-B**).

To experimentally test whether these residues are involved in hIRE1*α* LD’s interaction with peptides, we expressed and purified these mutants and conducted fluorescence anisotropy experiments using fluorescently labeled MPZ1N-2X peptide. We could purify both E102R and Y161R mutants to high purity (**Supplementary Fig. 18C**). They both behaved similarly to the wild type during purification. Notably, both E102R and Y161R mutants demonstrated around two-fold lower binding affinity (**Fig. 4G**, E102 *K*_1*/*2_= 6.35 *μM* and Y161R *K*_1*/*2_= 5.4 *μM*, **Supplementary Table 3**) compared to the wild-type (*K*_1*/*2_= 2.14 *μM*, **Supplementary Table 3**), revealing that the protein’s central area is crucial for binding unfolded proteins and that binding activity occurs within the pocket defined by E102 and Y161.

In summary, the E102R and Y161R mutations of cLD destabilized the binding of MPZ1N-2X by lowering the interactions at the center of the hIRE1*α* cLD dimer.

### hIRE1*α* cLD intermolecular interactions guide the activation process

Throughout the activation process, the hIRE1*α* cLD forms multiple intermolecular interactions, engaging not only with unfolded proteins but also with other cLDs and BiP. Karagöz et al. proposed a direct activation model whereby unfolded proteins facilitate IRE1 clustering by simultaneously engaging multiple dimers.^23^ Here, we examined whether our understanding of cLD-unfolded polypeptide interactions would be consistent with this model. We conducted simulations of a system with two cLD dimers bridged by one copy of MPZ1N-2X (**Fig. 5A**). The complex was established following the same principles as for the single dimer, and the initial configuration was optimized to position the peptide residues near critical residues identified in the contact site analysis, particularly Y161. Throughout the 200 ns of simulation, the two dimers were stably bridged by the peptide. This simulation offers a structural model for the role of unfolded proteins during the initial phase of cluster formation, where it promotes the accumulation of hIRE1*α*.

**Figure 5.**
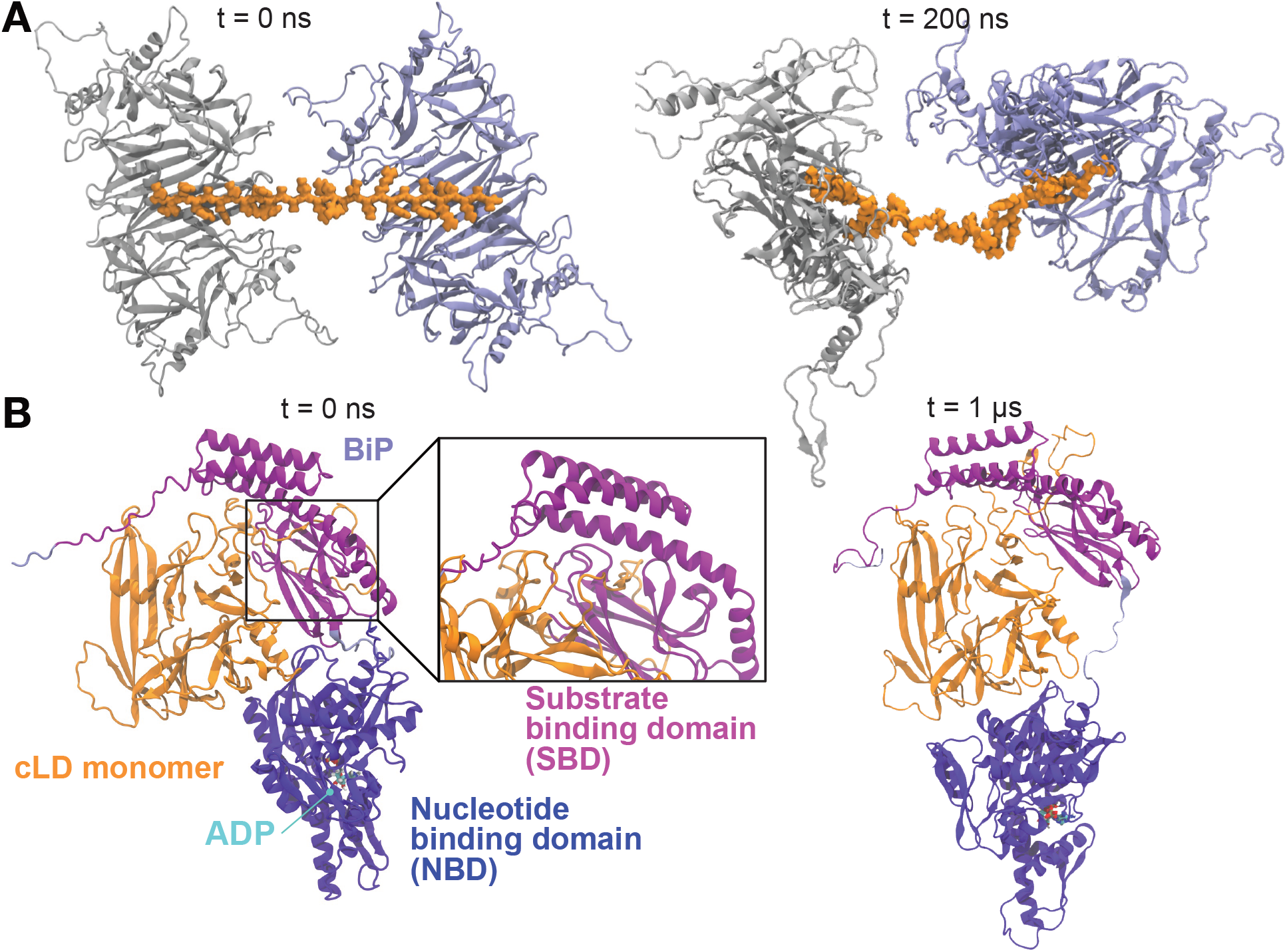
hIRE1*α* cLD intermolecular interactions. (A) Model of two cLD dimers interacting via an unfolded peptide (MPZ1N-2X): the system setup is shown (t = 0 ns) and a snapshot of the simulation (t = 200 ns). (B) BiP-cLD monomer complex as predicted by AlphaFold (BiP in shades of purple, cLD in orange) before the simulation (t = 0 *μ*s) and at the end of the simulation (t = 1 *μ*s). The SBD (residues E19-D408) is colored in light purple, and the NDB (residues C420-E650) in dark purple, and the interdomain linker (residues D409-V419) and KDEL motif (residues K651-L654) in light purple.

We used AlphaFold 3 to model the interaction between a cLD monomer and BiP (residues E19–L654) in the presence of ATP and ADP (**Fig. 5B, Supplementary Fig. 19A**). Prediction quality was limited in the apo and ADP-bound states (pTM = 0.48, ipTM = 0.59; pTM = 0.49, ipTM = 0.61, respectively), whereas ATP binding improved accuracy (pTM = 0.66, ipTM = 0.72). The predicted interfaces involved DR2, particularly residues 314-PLLEG-318, forming a short parallel *β*-sheet with the substrate-binding domain (SBD) of BiP through two hydrogen bonds. All AlphaFold 3 models were stable across three 1-μs simulations (**Supplementary Fig. 19B**), with cLD–BiP interfaces retaining 60–80% of initial contacts (**Supplementary Fig. 20**). In the apo and ADP-bound states, the nucleotide-binding domain (NBD) showed high Predicted Aligned Error (PAE) relative to the cLD, indicating uncertain positioning of the two domains relative to each other. Notably, in the ADP-bound state, which is thought to interact with hIRE1*α* cLD, the NBD remained mobile but proximal to the *α*B-helices, thereby restricting access to this region. Together, the AlphaFold 3 predictions suggest that BiP engages hIRE1*α* cLD by sterically hindering the oligomerization interface defined by DR2 and the *α*B-helices.^14^

## Discussion

IRE1 is the most conserved activator of the UPR across all eukaryotes.^6,7^ It responds to unfolded protein accumulation in the ER. Understanding the early activation stages of human IRE1*α* presents significant challenges due to conflicting evidence regarding the mechanism of IRE1’s interaction with unfolded peptides.^14,25^ To clarify these early activation stages, we conducted extensive atomistic MD simulations of the human IRE1*α* cLD and its interaction partners.

Our simulations revealed that the human IRE1*α* cLD forms stable dimers in solution under conditions similar to *in vitro* experiments and non-stress situations. Our findings on dimer stability support the recent observation that IRE1 is present in cells as a constitutive homodimer^29^ that forms oligomers in case of ER stress. We used two distinct models of the cLD dimer: one derived from X-ray crystallography^24^ (the “PDB model”) and the other predicted by AlphaFold 2 Multimer (the “AF model”). ^33,34^ The comparison between the two models was fundamental for identifying key structural elements. During the modeling of the missing disordered regions on the PDB model, we identified Y166 and L116 side chain orientation as crucial for the stability of the dimer. In fact, some mutants in this region that disturb the formation of dimers were previously identified by size exclusion chromatography, ^24,25^ including W125A, which is in contact with the residues we identified. Furthermore, we leveraged available data on HDX-MS experiments^25^ to confirm that the dimeric interface is stable and observe that the *α*B-helices might be shorter than what is observed in the crystallographic structure. This is an interesting observation, given that the *α*B-helices are supposed to form the oligomerization interface of human IRE1*α* cLD.^14^ Comparing simulation results to experimental data helped select the optimal dimer model and water force field, which is crucial for accurately reproducing the ensemble of disordered regions.

Credle et al. proposed that in *S. cerevisiae*, the Ire1 cLD dimer interacts with unfolded proteins through a mechanism similar to the peptide-binding mode of MHC-I, where peptides occupy a central cleft.^12^ Our simulations indicate that in comparison to the yeast IRE1 cLD dimer, the structure of the human dimer has a narrower central groove and is also gated by two glutamine side chains. This conformation prevents the formation of an ‘open’ peptide-binding central groove.

We tested several unfolded polypeptides for binding to the human IRE1*α* cLD dimer and found that peptides enriched in positive and aromatic residues can stably bind to the central part of the dimer, thanks to negative, polar, and aromatic amino acids. However, the binding does not take place inside the central groove. We found that the residues involved in peptide binding belong to the *α*A-helices, particularly E102, and to two pockets per monomer, characterized by residues Y117 and Y161. Among the most important contacts, we also identified L186, which was the main source of signal in the paramagnetic relaxation enhancement (PRE) experiments performed by spin-labeled peptides. ^14^ We found that peptides are able to bind to a ‘closed’ groove conformation, which is in line with fluorescence polarization experiments, showing that peptides can still bind to IRE1 LD Q105C SS, where a disulfide bridge locks the dimer’s central groove.^25^

We found that the E120R and Y161R mutations destabilize the binding of MPZ1N-2X by reducing the interactions between the unfolded peptide and the central region of the hIRE1*α* cLD dimer. The residue E102 is an important hotspot for binding on the center of the dimer, and with its negative charge, can be a selection for binding positively charged peptides. The residue Y161 is situated near L186, which is recognized as a critical hotspot for peptide binding based on PRE experiments with hIRE1*α*.^14^ In addition, a mutation in the corresponding residue in yIre1 negatively affects protein activation. ^12^ These findings were corroborated by fluorescence anisotropy experiments, which showed that Y161R and E102R mutants display lower binding affinity for the MPZ1N-2X compared to the wild-type hIRE1*α* LD. Altogether, our data suggest that the pocket defined by E102 and Y161 mediates peptide binding.

We conducted further exploration of the intermolecular interactions involved in the early stages of hIRE1*α* activation. During this process, the protein interacts with unfolded proteins, other hIRE1*α* monomers and dimers, and BiP. Unfolded proteins may serve as linkers between cLD dimers under ER stress,^23^ and Kettel et al. demonstrated that short peptides can facilitate clustering.^26^ Based on the knowledge acquired here on key contact sites, we constructed a system where the MPZ1N-2X peptide could maintain a stable interaction with two dimers at the same time. We speculate that unfolded proteins may bring dimers together in the early stage of ER stress and cluster formation, establishing assemblies with undefined stoichiometry and conformation. Structured oligomers, as observed in crystals by Credle et al.^12^ and *in situ* by Tran et al.,^11^ might arise at later stages after allosteric rearrangement.^14^ On the other hand, we showed that a complex formed by cLD and BiP, as predicted by AlphaFold, is stable in equilibrium simulations. These findings are in line with the work of Amin-Wetzel et al. and Dawes et al., ^25,46^ who implied that DR2 should be involved in interactions with BiP. Kettel et al.^26^ showed that mutating residues ^312^TLPL^315^, which partially overlap with the predicted interaction interface, is detrimental to the formation of condensates and higher-order oligomers. These results hint at the role of BiP in disrupting the oligomers by shielding the DR2 and *α*B-helices, effectively blocking the oligomeric interface. We note that AlphaFold 3 distinguished between ligand-bound states by capturing open and closed conformations of the SBD lid, even though with limited prediction quality.^47^

Recently, Simpson et al. identified a new binding site for cholera toxin on hIRE1*α* cLD,^28^ which competes with C-terminal residues I362-P368. Here, however, we propose a different peptide binding mechanism that does not involve disordered regions, which could be a mode of action of a different set of peptides.^26^ Indeed, the motif identified by Simpson et al. (YGWYXXH) is not recognizable in the model peptides studied here.

Our equilibrium atomistic molecular dynamics simulations are unbiased and provide a high level of detail. However, time scale limitations restrict our ability to observe the slower conformational rearrangements of the protein complexes analyzed here. We employed a force field designed for disordered and unfolded proteins,^38^ but some bias could still exist. This could potentially lead to an overestimation of protein-protein interactions and hinder our observation of peptide dissociation in cases of intermediate binding affinities.

While here we focused on the activation of the UPR by an excessive accumulation of unfolded proteins, Ire1’s activation is also triggered by a perturbation of the membrane composition of the ER.^48–50^ Future work will focus on investigating a structural model that explains how Ire1 can integrate molecular stress signals both from the ER lumen and its membrane.

In summary, our research endorses a model for the activation of human IRE1*α*, where pre-existing dimers in the ER lumen directly engage with accumulating unfolded proteins via the cLD. These interactions could promote dimer aggregation and initiate the formation of larger supra-molecular assemblies. Furthermore, our structural data suggest that BiP may inhibit IRE1 from forming oligomers while allowing dimer presence. Therefore, the dissociation of hIRE1*α* dimers from BiP, followed by their direct engagement with unfolded proteins, might facilitate the nucleation of assemblies that act as precursors to organized oligomeric structures (**Fig. 6**). In conclusion, our results provide a comprehensive framework to understand the structural dynamics of the Ire1 activation.

**Figure 6.**
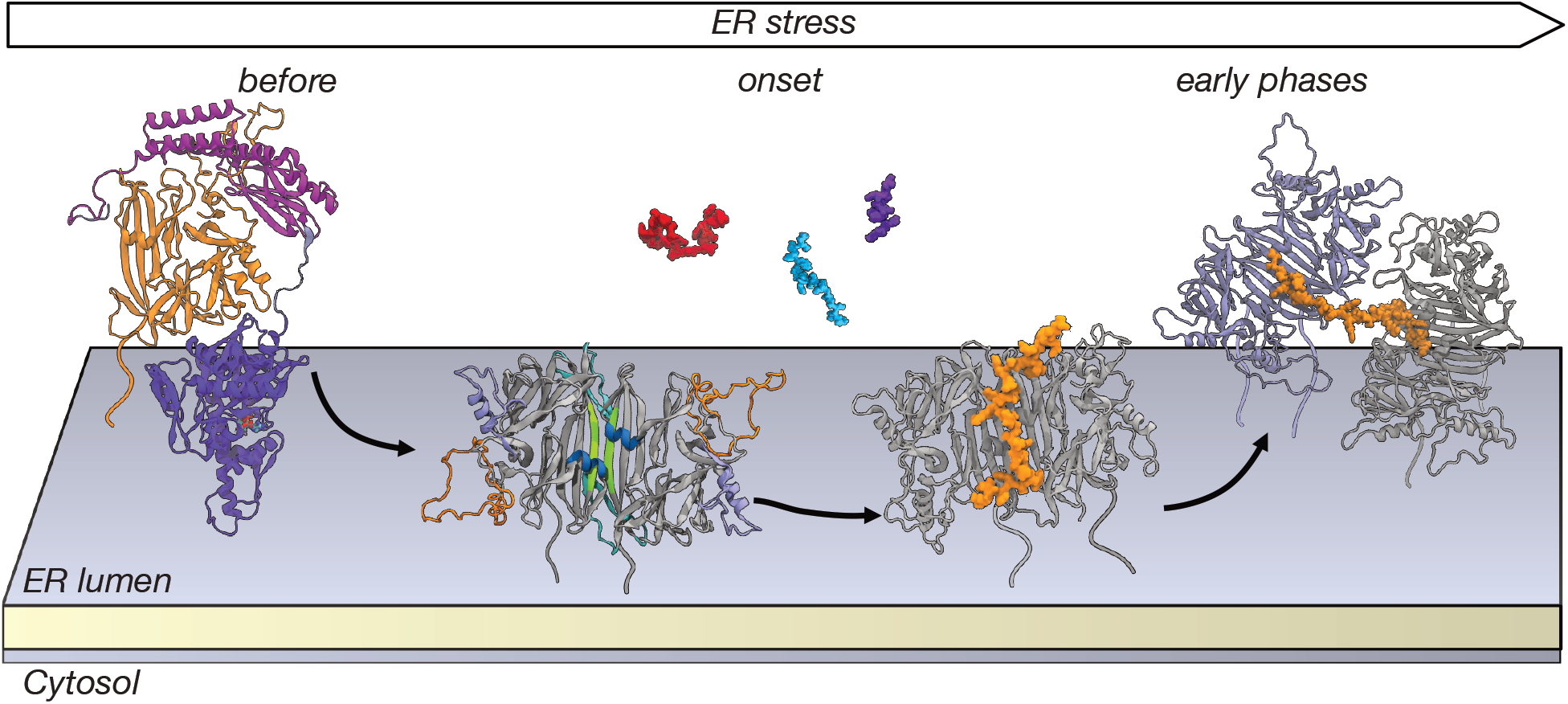
Schematic illustrating the proposed sequence of events occurring in the early stages of hIRE1*α* activation within the ER lumen. Before the onset of ER stress conditions, hIRE1*α* is prevented from forming assemblies by the chaperone BiP, which occupies the cLD’s oligomerization interface. As ER stress begins, unfolded proteins start to accumulate in the ER lumen, and pre-formed hIRE1*α* cLD dimers are released from BiP. Now, hIRE1*α* can interact with unfolded proteins, which brings multiple copies of the protein together to allow the formation of larger assemblies.

## Methods

### MD simulations

We performed all atomistic molecular dynamics simulations using GROMACS 2021.4 and 2024.3^51^ and the Charmm36m force field. ^41^ All systems were set up with explicit solvent, TIP3P^40^ or TIP4P-D^38^ water model, and 0.15 M NaCl, adding to the number of ions needed for a neutral total net charge of the system. The TIP4P-D water model was taken from the work of Piana et al.^38^ and inserted into the Charmm36m force field parameters by setting the parameters for the oxygen atom OWT4PD Lennard-Jones interactions to *<* = 0.9365543 kJ/mol and *σ* = 0.316499897100 nm and using a modified itp file of the TIP4P/2005 water model^52^ (available at http://catalan.quim.ucm.es/), setting the charge of H atoms to 0.58 and the charge of the dummy atom to −1.16. The solvation in TIP3P was done using CHARMM-GUI,^53^ while the solvation in TIP4P-D was done using GROMACS tools and inhouse bash scripts. The systems were minimized using the steepest descent algorithm and equilibrated in the NVT ensemble with position restraints (on the backbone and side chains) for 0.125 ns with a 1 fs time step at 300 K. The production runs were performed in the NPT ensemble. The systems were maintained at a temperature of 300 K using the velocity rescale thermostat^54^ (*τ*_*T*_ = 1 ps) and at 1 bar using the Parrinello-Rahman barostat^55^ (*τ*_*P*_ = 5 ps). The simulations performed in this work are summarised in the Table 1.

**Table 1:**
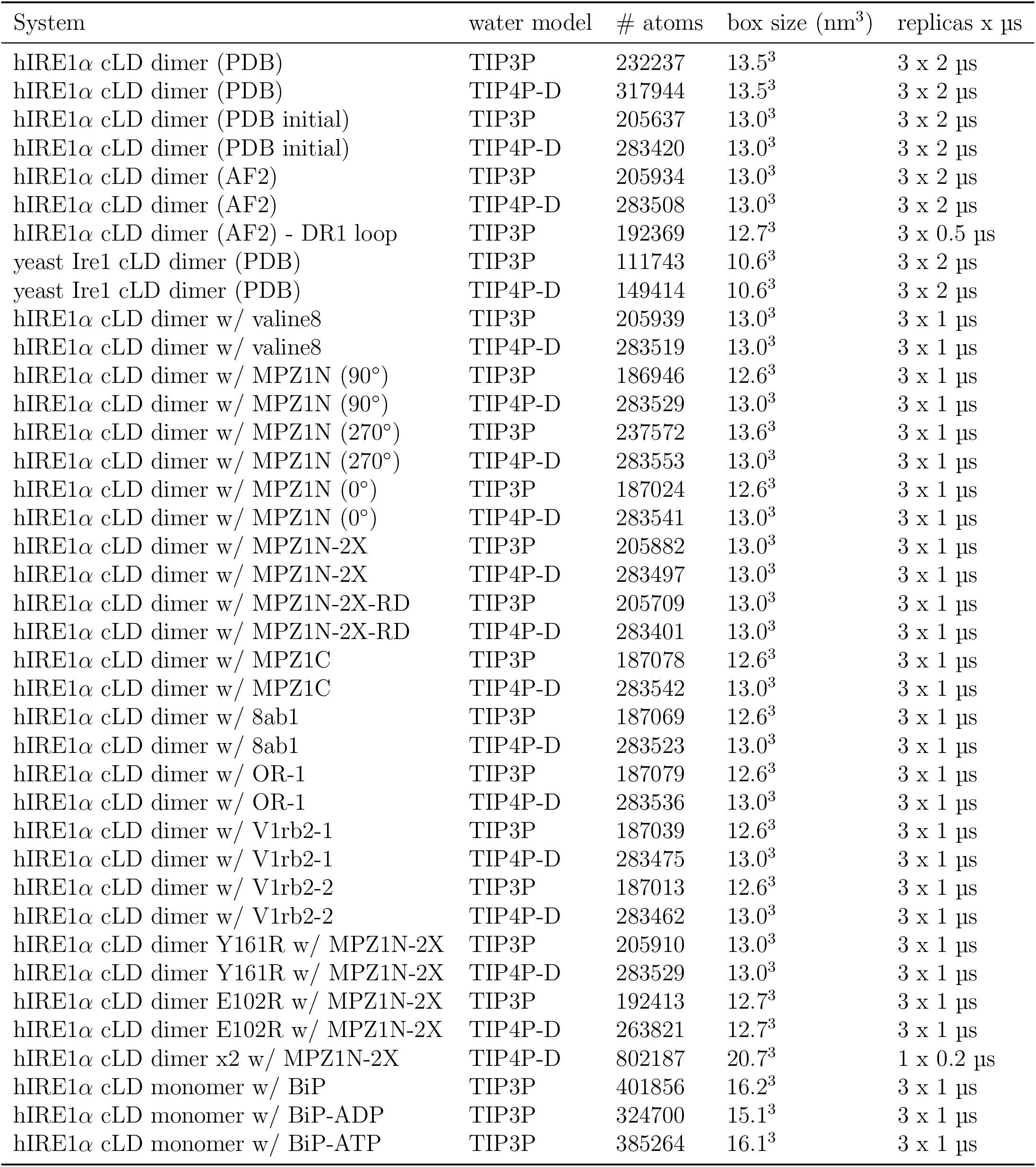
The simulations performed in this work are summarised here. All the peptides were simulated in complex with hIRE1*α* cLD dimer AlphaFold2 (AF2) model. All systems were solvated in either TIP3P or TIP4P-D water and with 0.15 M NaCl plus counterions. All simulation frames were saved with a timestep of 0.25 ns.

| System | water model | # atoms | box size (nm<sup>3</sup>) | replicas x μs |
| --- | --- | --- | --- | --- |
| hIRE1α cLD dimer (PDB) | TIP3P | 232237 | 13.5<sup>3</sup> | 3 x 2 μs |
| hIRE1α cLD dimer (PDB) | TIP4P-D | 317944 | 13.5<sup>3</sup> | 3 x 2 μs |
| hIRE1α cLD dimer (PDB initial) | TIP3P | 205637 | 13.0<sup>3</sup> | 3 x 2 μs |
| hIRE1α cLD dimer (PDB initial) | TIP4P-D | 283420 | 13.0<sup>3</sup> | 3 x 2 μs |
| hIRE1α cLD dimer (AF2) | TIP3P | 205934 | 13.0<sup>3</sup> | 3 x 2 μs |
| hIRE1α cLD dimer (AF2) | TIP4P-D | 283508 | 13.0<sup>3</sup> | 3 x 2 μs |
| hIRE1α cLD dimer (AF2) - DR1 loop | TIP3P | 192369 | 12.7<sup>3</sup> | 3 x 0.5 μs |
| yeast Ire1 cLD dimer (PDB) | TIP3P | 111743 | 10.6<sup>3</sup> | 3 x 2 μs |
| yeast Ire1 cLD dimer (PDB) | TIP4P-D | 149414 | 10.6<sup>3</sup> | 3 x 2 μs |
| hIRE1α cLD dimer w/ valine8 | TIP3P | 205939 | 13.0<sup>3</sup> | 3 x 1 μs |
| hIRE1α cLD dimer w/ valine8 | TIP4P-D | 283519 | 13.0<sup>3</sup> | 3 x 1 μs |
| hIRE1α cLD dimer w/ MPZ1N (90°) | TIP3P | 186946 | 12.6<sup>3</sup> | 3 x 1 μs |
| hIRE1α cLD dimer w/ MPZ1N (90°) | TIP4P-D | 283529 | 13.0<sup>3</sup> | 3 x 1 μs |
| hIRE1α cLD dimer w/ MPZ1N (270°) | TIP3P | 237572 | 13.6<sup>3</sup> | 3 x 1 μs |
| hIRE1α cLD dimer w/ MPZ1N (270°) | TIP4P-D | 283553 | 13.0<sup>3</sup> | 3 x 1 μs |
| hIRE1α cLD dimer w/ MPZ1N (0°) | TIP3P | 187024 | 12.6<sup>3</sup> | 3 x 1 μs |
| hIRE1α cLD dimer w/ MPZ1N (0°) | TIP4P-D | 283541 | 13.0<sup>3</sup> | 3 x 1 μs |
| hIRE1α cLD dimer w/ MPZ1N-2X | TIP3P | 205882 | 13.0<sup>3</sup> | 3 x 1 μs |
| hIRE1α cLD dimer w/ MPZ1N-2X | TIP4P-D | 283497 | 13.0<sup>3</sup> | 3 x 1 μs |
| hIRE1α cLD dimer w/ MPZ1N-2X-RD | TIP3P | 205709 | 13.0<sup>3</sup> | 3 x 1 μs |
| hIRE1α cLD dimer w/ MPZ1N-2X-RD | TIP4P-D | 283401 | 13.0<sup>3</sup> | 3 x 1 μs |
| hIRE1α cLD dimer w/ MPZ1C | TIP3P | 187078 | 12.6<sup>3</sup> | 3 x 1 μs |
| hIRE1α cLD dimer w/ MPZ1C | TIP4P-D | 283542 | 13.0<sup>3</sup> | 3 x 1 μs |
| hIRE1α cLD dimer w/ 8ab1 | TIP3P | 187069 | 12.6<sup>3</sup> | 3 x 1 μs |
| hIRE1α cLD dimer w/ 8ab1 | TIP4P-D | 283523 | 13.0<sup>3</sup> | 3 x 1 μs |
| hIRE1α cLD dimer w/ OR-1 | TIP3P | 187079 | 12.6<sup>3</sup> | 3 x 1 μs |
| hIRE1α cLD dimer w/ OR-1 | TIP4P-D | 283536 | 13.0<sup>3</sup> | 3 x 1 μs |
| hIRE1α cLD dimer w/ V1rb2-1 | TIP3P | 187039 | 12.6<sup>3</sup> | 3 x 1 μs |
| hIRE1α cLD dimer w/ V1rb2-1 | TIP4P-D | 283475 | 13.0<sup>3</sup> | 3 x 1 μs |
| hIRE1α cLD dimer w/ V1rb2-2 | TIP3P | 187013 | 12.6<sup>3</sup> | 3 x 1 μs |
| hIRE1α cLD dimer w/ V1rb2-2 | TIP4P-D | 283462 | 13.0<sup>3</sup> | 3 x 1 μs |
| hIRE1α cLD dimer Y161R w/ MPZ1N-2X | TIP3P | 205910 | 13.0<sup>3</sup> | 3 x 1 μs |
| hIRE1α cLD dimer Y161R w/ MPZ1N-2X | TIP4P-D | 283529 | 13.0<sup>3</sup> | 3 x 1 μs |
| hIRE1α cLD dimer E102R w/ MPZ1N-2X | TIP3P | 192413 | 12.7<sup>3</sup> | 3 x 1 μs |
| hIRE1α cLD dimer E102R w/ MPZ1N-2X | TIP4P-D | 263821 | 12.7<sup>3</sup> | 3 x 1 μs |
| hIRE1α cLD dimer x2 w/ MPZ1N-2X | TIP4P-D | 802187 | 20.7<sup>3</sup> | 1 x 0.2 μs |
| hIRE1α cLD monomer w/ BiP | TIP3P | 401856 | 16.2<sup>3</sup> | 3 x 1 μs |
| hIRE1α cLD monomer w/ BiP-ADP | TIP3P | 324700 | 15.1<sup>3</sup> | 3 x 1 μs |
| hIRE1α cLD monomer w/ BiP-ATP | TIP3P | 385264 | 16.1<sup>3</sup> | 3 x 1 μs |

**Table 2:** Available binding affinities (K_1/2_) of the unfolded polypeptides studied to hIRE1*α*.^14,26^.

| Polypeptide | $K_{1/2}[\mu M]$ |
| --- | --- |
| MPZ1-N | 16 |
| MPZ1N-2X | 0.456 |
| MPZ1-C | 572 |
| MPZ1N-2X-RD | binding impaired |
| 8ab1 | 5 |

**Table 3:** Goodness-of-fit parameters of binding curves of MPZ1-N-2X peptide to hIRE1*α* LD for fluorescence anisotropy experiments.

| | hIRE1α LD wild-type | hIRE1α LD E102R | hIRE1α LD Y161R |
| --- | --- | --- | --- |
| $K_D$ [$\mu M$] | 2.14 | 6.35 | 5.433 |
| $K_D$ 95% confidence interval [$\mu M$] | 1.809 to 2.528 | 5.835 to 6.959 | 4.296 to 7.095 |
| $R^2$ | 0.9918 | 0.9992 | 0.9851 |

### IRE1*α* core Luminal Domain (cLD) structural models

#### Human PDB dimer

The protein structure of the human IRE1*α* core Luminal Domain (cLD) dimer was obtained from the Protein Data Bank, PDB ID 2HZ6.^24^ The missing residues were modeled using Modeller in UCSF Chimera^56^ (version 1.15 and 1.17), and they included disordered regions 1 and 2 in residues 131-152 and 308-357 and three connecting loops in residues 66-70, 89-90, and 11-115.

An initial PDB model was briefly equilibrated in NPT, and a conformation with a groove width of approximately 0.6 nm was selected. This snapshot was used as the initial structure for the initial “PDB model” simulations, in which the dimer dissociates.

The final PDB model was obtained from the deposited crystal structure, adding missing residues with Modeller in UCSF Chimera independently for the two monomers and not allowing residues at the borders of the missing areas to rearrange during modeling. This model was then directly used for simulation setup, producing the stable cLD dimer used here.

#### Human AlphaFold dimer

We obtained the AlphaFold prediction of the human IRE1*α* cLD dimer (“AF model”) by running AlphaFold 2 Multimer^33,34^ in the ColabFold^36^ notebook v1.5.5 using MMseq2 (https://colab.research.google.com/github/sokrypton/ColabFold/blob/main/AlphaFold2.ipynb) with disabled template usage.

The structures of the five top models were predicted, and the first one was chosen for simulation. The input sequence was (Uniprot identifier O75460): PETLLFVSTLDGSLHA VSKRTGSIKWTLKEDPVLQVPTHVEEPAFLPDPNDGSLYTLGSKNNEGLTKLPFTIPELVQASPCRSSDGILYMGKKQDIWYVIDLLTGEKQQTLSSAFADSLCPSTSLLYLGRTEYTITMYDTKTRELRWNATYFDYAASLPEDDVDYKMSHFVSNGDGLVVTVDSESGDVLWIQNYASPVVAFYVWQREGLRKVMHINVAVETLRYLTFMSGEVGRITKWKYPFPKETEAKSKLTPTLYVGKYSTSLYASPSMVHEGVAVVPRGSTLPLLEGPQTDGVTIGDKGECVITPSTDVKFDPGLKSKNKLNYLRNYWLLIGHHETP.

For the prediction of the AlphaFold 3 model of human IRE1*α* cLD dimer we used the AlphaFold server^57^ at https://alphafoldserver.com/ with same input sequence as for AlphaFold 2.

#### Yeast PDB dimer

The protein structure of the yeast Ire1 Luminal Domain dimer was obtained from the Protein Data Bank entry 2BE1.^12^ The system was set up using CHARMM-GUI, which was also used to add missing residues (210-219, 255-274, 380-387). The model simulated here spans residue 111 to 449 (Uniprot identifier P32361).

#### DR1-loop AlphaFold dimer

The model of the human IRE1*α* cLD dimer predicted by the AlphaFold multimer was modified by removing the DR1 (residues 131-152) and adding them again as unstructured loops, using Modeller in UCSF Chimera^56^ (version 1.17.3).

### Human IRE1*α* cLD in complexes

#### Unfolded peptides

For the binding experiments, we created an unfolded and stretched structure from the sequence of interest using the UCSF Chimera ‘Build structure’ tool. The peptides we simulated mimicked unfolded peptides, so no specific secondary structure was imposed. The stretched-out configuration was obtained by setting the seed for ϕ/*ψ* dihedrals with values for antiparallel *β* strand to obtain angles ϕ = −139 and *ψ* = 135 for all the residues. Then, we manually placed the peptides above the binding groove of the AF model of human IRE1*α* cLD dimer. The sequences of the peptides used are: valine8, VVVVVVVV; MPZ1-N, LIRYAWLRRQAA; MPZ1N-2X, LIRYAWLRRQAALQRRLIRYAWLRRQAA; MPZ1N-2X-RD, LIDYAWLDDQAALQDDLIDYAWLDDQAA; MPZ1-C, LQRRISAME; 8ab1, WLCALGKVLPFHRWHTMV; OR-1, MEKAVLINQTSVMSFR; V1rb2-1, MFMPWGRWNSTTC-QSLIYLHR; V1rb2-2, LKFKDCSVFYFVHIIMSHSYA.

For the prediction of the AlphaFold 3 model of human IRE1*α* cLD dimer in complex with MPZ1N-2X, we used the AlphaFold server^57^ at https://alphafoldserver.com/ with same input sequences as for AlphaFold 2.

#### E102R and Y161R AlphaFold dimer and MPZ1N-2X peptide

We created the mutants E102R and Y161R of human IRE1*α* cLD dimer by substituting the tyrosine residue at position 161 on both monomers with arginines in the AlphaFold dimer model in complex with the unfolded peptide MPZ1N-2X, utilizing the mutation options in CHARMM-GUI,^53^ during the setup of the system.

#### Two dimers and MPZ1N-2X peptide

The system, which contains two copies of the AF model of the human IRE1*α* cLD dimer and one copy of the unfolded peptide MPZ1N-2X, was obtained by manually arranging the copies in UCSF Chimera.^56^

#### cLD monomer in complex with BiP

The BiP-cLD heterodimer systems were predicted with AlphaFold 3 using the AlphaFold server^57^ at https://alphafoldserver.com/. The hIRE1*α* cLD sequence used is the same used for predicting the dimer: the PDB 2HZ6 sequence, Uniprot identifier O75460 with mutations C127S and C311S, and residues P29-P368. The BiP sequence used is taken from UniProt identifier P11021, residues E19-L654. We predicted three complexes: one without any nucleotide, one containing ADP, and another containing ATP. Simulations of the BiP-cLD complex were run in TIP3P water.

### Hydrogen-deuterium exchange fractions calculation from MD simulations

We based our HDX-MS data analysis on the method developed by Bradshaw et al. ^43^ to compare experimental HDX-MS data to simulations. We exploited the first part of the pipeline (script ‘calc hdx.py’) to compare our simulation results for the trajectories concerning the cLD dimer to the experimental data from Amin-Wetzel et al.^25^

To determine the deuterated fraction of a peptide segment from simulations, the protection factor for each residue *i, P*_*i*_, must be computed from the simulation snapshots, following the approach of Best and Vendruscolo:^58^ *lnP*_*i*_ = ⟨
*β*_*C*_*N*_*C,i*_ + *β*_*H*_*N*_*H,i*_⟩. Here, *N*_*C,i*_ and *N*_*H,i*_ are the number of H-bonds and heavy-atom contacts of the backbone amide of residue *i*, and the scaling factors *β*_*C*_ and *β*_*H*_ are set to 0.35 and 2.0, respectively. The simulated deuterated fraction of a peptide segment, 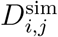, defined by residues *m*_*j*_ + 1 to *n*_*j*_, was then calculated at any exchange time point *t* as:

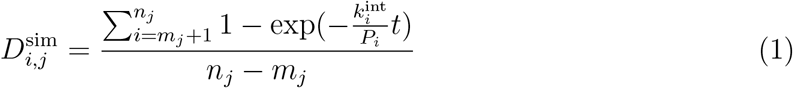

where *m*_*j*_ and *n*_*j*_ are the first and last residue numbers of the j-th protein fragment, respectively. The intrinsic exchange rate constants for each residue type 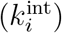 were obtained from Bai et al. with updated acidic residues and glycine.^59,60^

To reproduce the time points after incubation in deuterium (D_2_O), we computed deuterated fractions separately for each of the two monomers constituting a dimer for the time points 0.5 min (30 s) and 5 min (300 s). Then, we computed the mean and standard deviation over the data coming from replicas of the same cLD dimer model (AF or PDB model) and the same water model (TIP3P or TIP4P-D). To estimate the uncertainty of the mean values obtained from our datasets and the dataset from Amin-Wetzel et al. (^25^ Figure 3—source data 1), we applied a non-parametric bootstrap resampling procedure. For each sequence range from HDX-MS analysis, we treated the measurements from the N=6 independent datasets as independent samples, accounting for 3 replicas each with two monomers (6 monomers total). We then generated 10,000 bootstrap replicates by sampling the datasets with replacement, maintaining the same number of samples N in each resample. For each replicate, we calculated the mean at each sequence position. The resulting distribution of bootstrap means was used to compute the standard deviation as an estimate of the standard error. We computed the difference between simulation and experimental data (deuterated fraction discrepancy), and for each residue, we selected as the ‘best structure’ the model with the discrepancy closest to zero among PDB-TIP3P, PDB-TIP4P-D, AF-TIP3P, and AF-TIP4P-D systems.

### Groove width analysis

To monitor the groove width, we calculated the Euclidean distance between the C*α* atoms of residues Q105 belonging to the two monomers forming the hIRE1*α* cLD dimer at every frame of the trajectory. For the analysis of the yeast Ire1 cLD dimer groove, we computed the distance between the C*α* atoms of the residues S200 belonging to the two monomers (PDB ID 2BE1^12^).

### Groove-peptide distance analysis

We measured the groove-peptide distance as the minimum distance between any residue belonging to the helices flanking the central groove (residues P101-S107) and any residue of the unfolded peptide. To analyze the groove width and groove-peptide distances, we used MDAnalysis^61,62^ (version 2.3.0) and custom Python scripts.

### Analysis of cLD dimer-peptides contacts

We assessed the interactions between the cLD dimer and unfolded peptides throughout the simulations using the Python package MDTraj^63^ (version 1.9.5). A contact was deemed to occur when two heavy atoms (hydrogens were excluded from the analysis) were found at a distance of less than 0.45 nm. For each 10th frame of a simulation replica, a contact map for all heavy atoms was computed, with binary values for the presence or absence of a contact. These contact maps per frame were summed together and normalized by the number of frames to obtain the frequency of contacts for a given simulation replica. To obtain a ranking of the cLD amino acids in contact with unfolded peptides, the contact map was reduced to one dimension by summing all the contributions from all peptide residues and normalizing by the number of atoms in the peptide, resulting in an array matching the dimensions of the cLD dimer. Next, the 1-dimensional contact maps from all replicas were summed together and normalized by the number of replicas to obtain the final ‘contact score’ and ranking.

### Binding free energy calculations (MM/PBSA)

The binding free energy of noncovalently bound complexes of human IRE1 cLD and peptides was calculated with MM/PBSA (Molecular mechanics/Poisson-Boltzmann Surface Area) method via gmx MMPBSA (version 1.6.4).^64,65^ The Poisson-Boltzmann method was used to estimate the electrostatic contribution to solvation free energy as recommended for data obtained with the CHARMM force field. The contribution of the entropic term was omitted, obtaining effective binding free energy values, or enthalpy of binding (Δ*H*). We used the Single Trajectory Protocol (STP), using the cLD-peptide simulations as input. The calculations were performed on the last 250 ns of each replica. Single-term total non-polar solvation free energy (*inp* = 1) was used. The charmm radii (*PBRadii* = 7) was used to build amber topology files.^66^ The default parameters were applied for other terms.

### Protein purification

To express hIRE1*α* LD (24-443) human cDNA sequences were cloned into pET47b(+) to create a coding sequence with N-terminal His6-tag. Mutations of hIRE1*α* LD were introduced by overlap extension PCR and restriction cloning into pET47b(+). For expression of the proteins, the plasmid of interest was transformed into Escherichia coli strain BL21DE3* RIPL (Agilent Technologies). Cells were grown in Luria Broth until OD600=0.6-0.8. Protein expression was induced with 0.6 mM IPTG, and cells were grown in 20°C overnight. For purification, cells after harvesting were resuspended in Lysis Buffer (50 mM HEPES pH 7.2, 400 mM NaCl, 20 mM imidazole, 5% glycerol, 5 mM *β*-mercaptoethanol) and were lysed in Constans Systems cell disruptor at 25 000 psi. The supernatant was collected after centrifugation for 45 minutes at 48000×g in 4°C. Supernatant was loaded onto Ni-NTA column (Cytiva) and the protein eluted with a linear gradient of imidazole from 20 to 500 mM. Fractions containing the protein were diluted 1:8 with anion exchange wash buffer (50 mM HEPES pH 7.2, 5 mM *β*-mercaptoethanol), loaded onto HiTRAP-Q ion exchange column (Cytiva) and eluted with a linear gradient from 50 mM to 1 M NaCl. Afterwards, the His6-tag was removed by cleavage with Precission protease (GE Healthcare, 1 μg of enzyme per 100 μg of protein). The cleavage was performed overnight in 4°C. The protein sample after cleavage was loaded onto a Ni-NTA column, and the flow-through containing protein without the tag was collected. The protein was further purified on a Superdex 200 10/300 gel filtration column equilibrated with Buffer A (25 mM HEPES pH 7.2, 150 mM NaCl, 2 mM DTT). Protein concentrations were determined using extinction coefficient at 280 nm predicted by the Expasy ProtParam tool (http://web.expasy.org/protparam/).

### Fluorescence anisotropy

For fluorescence anisotropy measurements, the MPZ1-N-2X peptide attached to 5 carboxyfluorescein (5-FAM) at its N-terminus was obtained from GenScript at *>*95% purity. Binding affinities of hIRE1*α* LD mutants to FAM-labeled peptides were determined by measuring the change in fluorescence anisotropy on a Tecan CM Spark Micro Plate Reader with excitation at 485 nm and emission at 525 nm with increasing concentrations of hIRE1*α* LD variants. Measurements were performed in Buffer A supplemented with Tween 20 (25 mM HEPES pH 7.2, 150 mM NaCl, 2 mM DTT, 0.025% Tween 20). Fluorescently labeled peptides were used in a concentration of 90 nM. The reaction volume of each data point was 25 *μ*L and the measurements were performed in 384-well, black flat-bottomed plates (Corning) after incubation of peptide with hIRE1*α* LD variants for 30 min at 25 °C. Binding curves were fitted using Prism Software (GraphPad) using the following equation: *F*_*bound*_ = *r*_*free*_ + (*r*_*max*_ − *r*_*free*_)*/*(1 + 10((*LogK*_1*/*2_ − *x*) · *n*_*H*_)), where *F*_*bound*_ is the fraction of peptide bound, *r*_*max*_ and *r*_*free*_ are the anisotropy values at maximum and minimum plateaus, respectively. *n*_*H*_ is the Hill coefficient and *x* is the concentration of the protein in log scale. Curve-fitting was performed with minimal constraints to obtain *K*_1*/*2_ values with high *R*^2^ values. However, as this equation does not consider the equilibria between hIRE1*α* LD dimers/oligomers, these apparent *K*_1*/*2_ values do not reflect the dissociation constant.

## Data availability statement

All input files, analysis script, and data presented in this work are freely available at https://doi.org/10.5281/zenodo.17063387.

## Acknowledgement

We thank Jan Stuke for the helpful insights and discussions, and Prof. David Ron for critical feedback and discussions. E.S. and R.C. acknowledge the support of Goethe University Frankfurt, the Frankfurt Institute of Advanced Studies, the LOEWE Center for Multiscale Modelling in Life Sciences (CMMS) of the state of Hesse, and the CRC 1507: Membrane-associated Protein Assemblies, Machineries, and Supercomplexes (P09), as well as computational resources and support from the Center for Scientific Computing of the Goethe University and the Jülich Supercomputing Centre. This research was funded in whole or in part by the Austrian Science Fund (FWF) [FWF-W1261, FWF-SFB F79] to G.E.K.

## Author Contributions

Conceptualization: R.C. Investigation, data curation, formal analysis, visualization: E.S. (modelling and simulations), G.Ś. (experimental validation). Writing: E.S., G.E.K., R.C. Project administration: R.C. Supervision, funding acquisition: G.E.K., R.C.

## Supporting Information Available

The molecular dynamics data presented in this work are available at https://doi.org/10.5281/zenodo.17063387.

## Supplementary Figures and Tables

**Figure 7.**
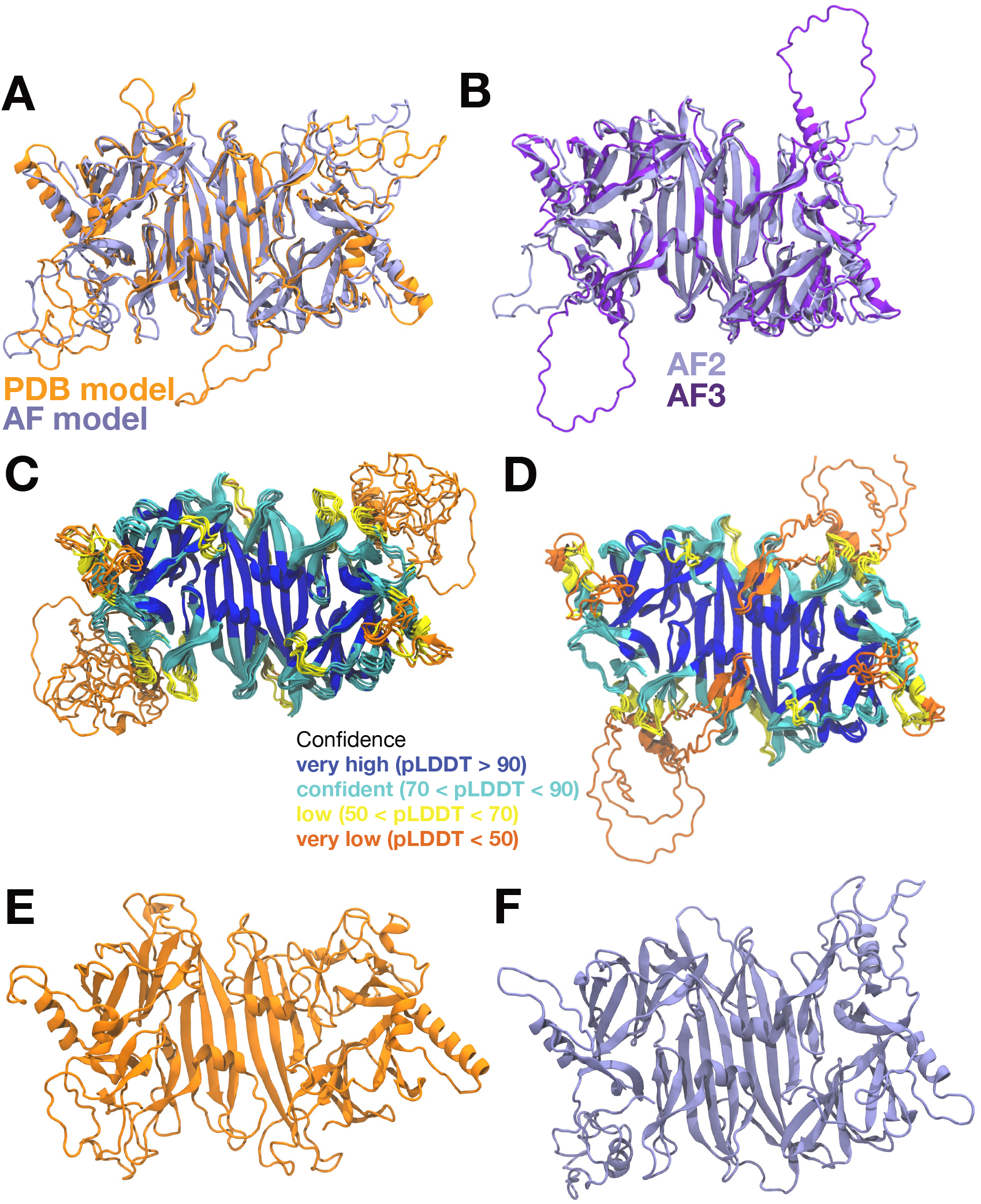
(A) Superposition of AF model (light violet) and PDB model (orange) of the hIRE1*α* cLD dimer before the simulation (RMSD = 3.34 Å). (B) Superposition of cLD dimer models predicted by AlphaFold 2 Multimer (light violet) and AlphaFold 3 (dark violet). (C) Superposition of the 5 structures predicted by AlphaFold 2 Multimer for the cLD dimer and colored by confidence prediction score (pLDDT). (D) Superposition of the 5 structures predicted by AlphaFold 3 for the cLD dimer and colored by confidence prediction score (pLDDT). (E) PDB model of the hIRE1*α* cLD dimer after 2 μs of production run from one of the TIP3P replicas. (F) AF model of the hIRE1*α* cLD dimer after 2 μs of production run from one of the TIP3P replicas.

**Figure 8.**
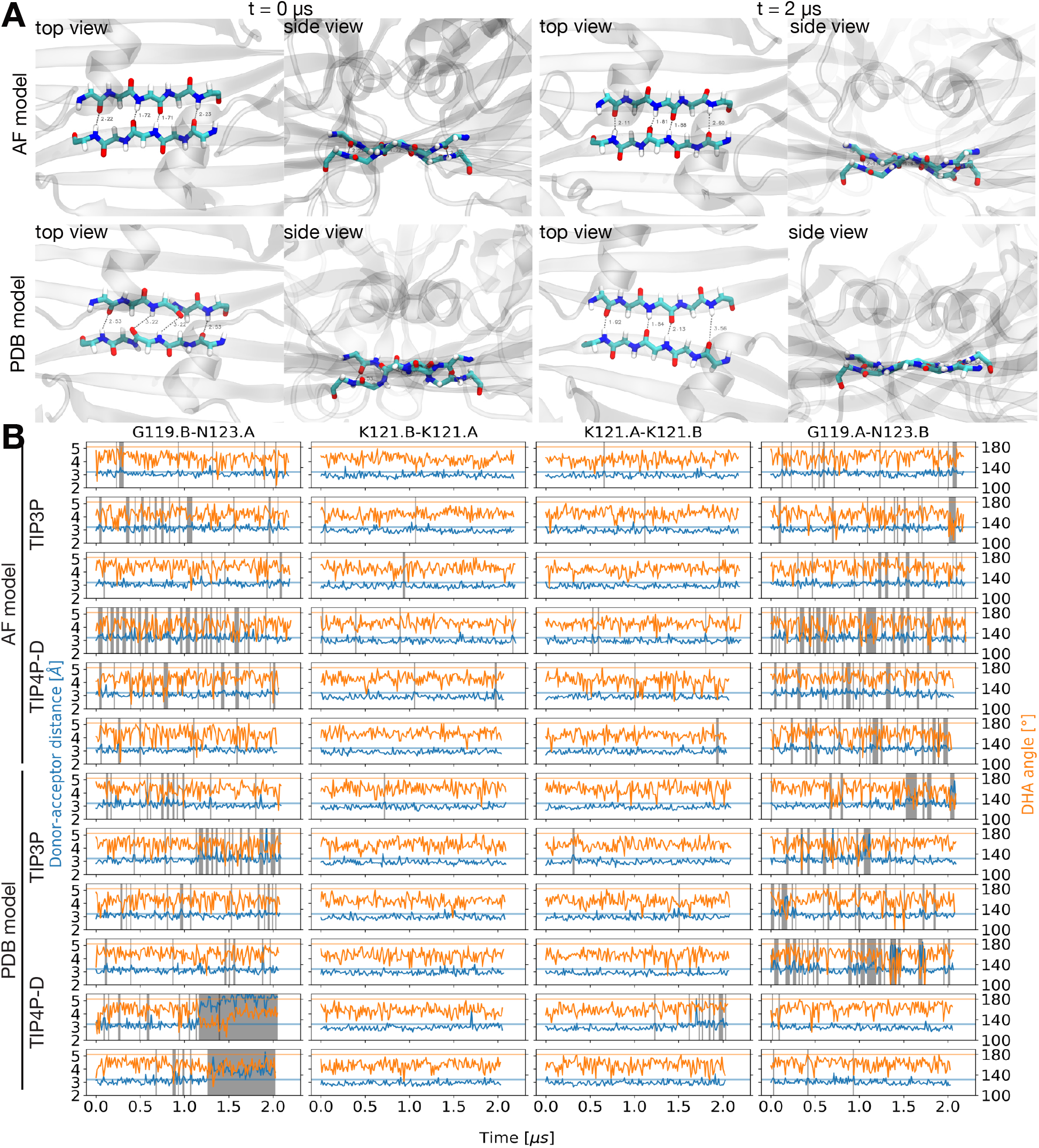
(A) Details of the dimeric interface of the hIRE*α* cLD dimer, consisting of an antiparallel *β*-sheet with four hydrogen bonds between residues G119-D123. The renders show the dimer interface in the AF model (upper row) and of the PDB model (lower row) prior to energy minimization and production run (t = 0 μs) and after 2 μs of simulation. The backbone atoms of residues G119-D123 are in sticks (C cyan, H white, O red, N blue) and the rest of the structure is rendered as a transparent ribbon. The backbone of the protein is shown as transparent cartoon. (B) Hydrogen bond analysis of all the hIRE*α* cLD dimer simulation replicas. This analysis shows the distance between donor and acceptor atoms (left y-axis, blue line) and the angle between donor-hydrogen-acceptor (right y-axis, orange line) for all replicas of PDB and AF models in TIP3P and TIP4P-D water. The hydrogen bonds analyzed here are those formed at the dimeric interface by residues G119, K121, and N123. The horizontal blue and orange lines indicate, respectively, the ideal distance (3.2 Å) and angle (180°) for a hydrogen bond. Whenever the distance is greater than 3.2 Å or the angle is lower than 120°, the frame is highlighted by a grey or red vertical bar, respectively. Therefore, the vertical bars indicate when the hydrogen bond is lost. The data are plotted every 50th frame for the lines and every frame for the vertical bars.

**Figure 9.**
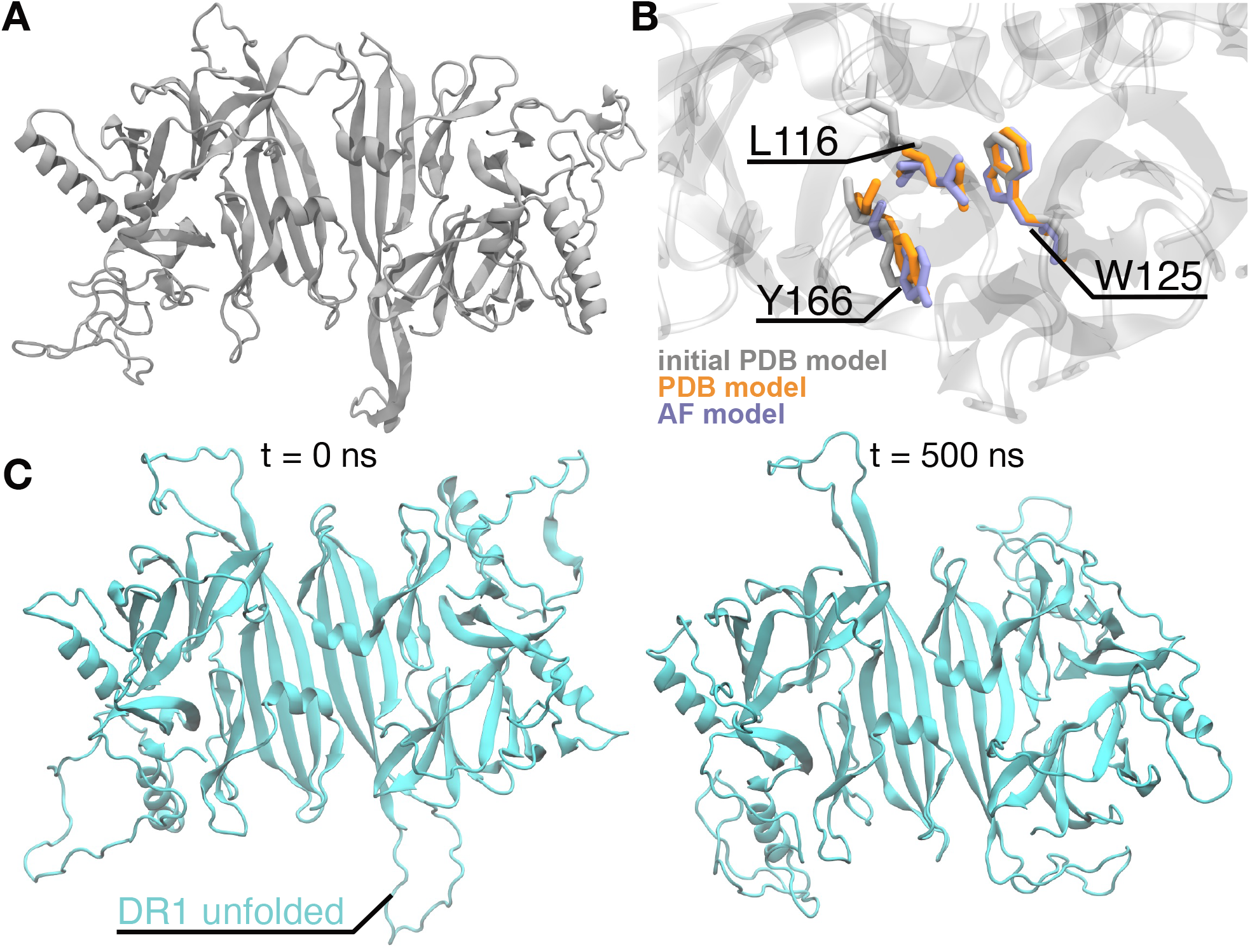
(A) Initial PDB model of the hIRE1*α* cLD dimer after 2 μs of production run. Replica showing a partial dissociation of the dimer. (B) Detail of the side of the dimeric interface of the initial PDB model that dissociates during simulations. The side chains of residues L116, Y166, and W125 are highlighted in three different models: the initial PDB model (gray), the final PDB model (orange), and the AlphaFold model (light violet). The backbone of the protein is shown as transparent cartoon. (C) The DR1 is folded in the AF model (Fig. 1B). The modified AF model with unfolded DR1 (left, t = 0 μs). The AlphaFold model with unfolded DR1 is stable after 500 ns of simulation (right, t= 500 μs).

**Figure 10.**
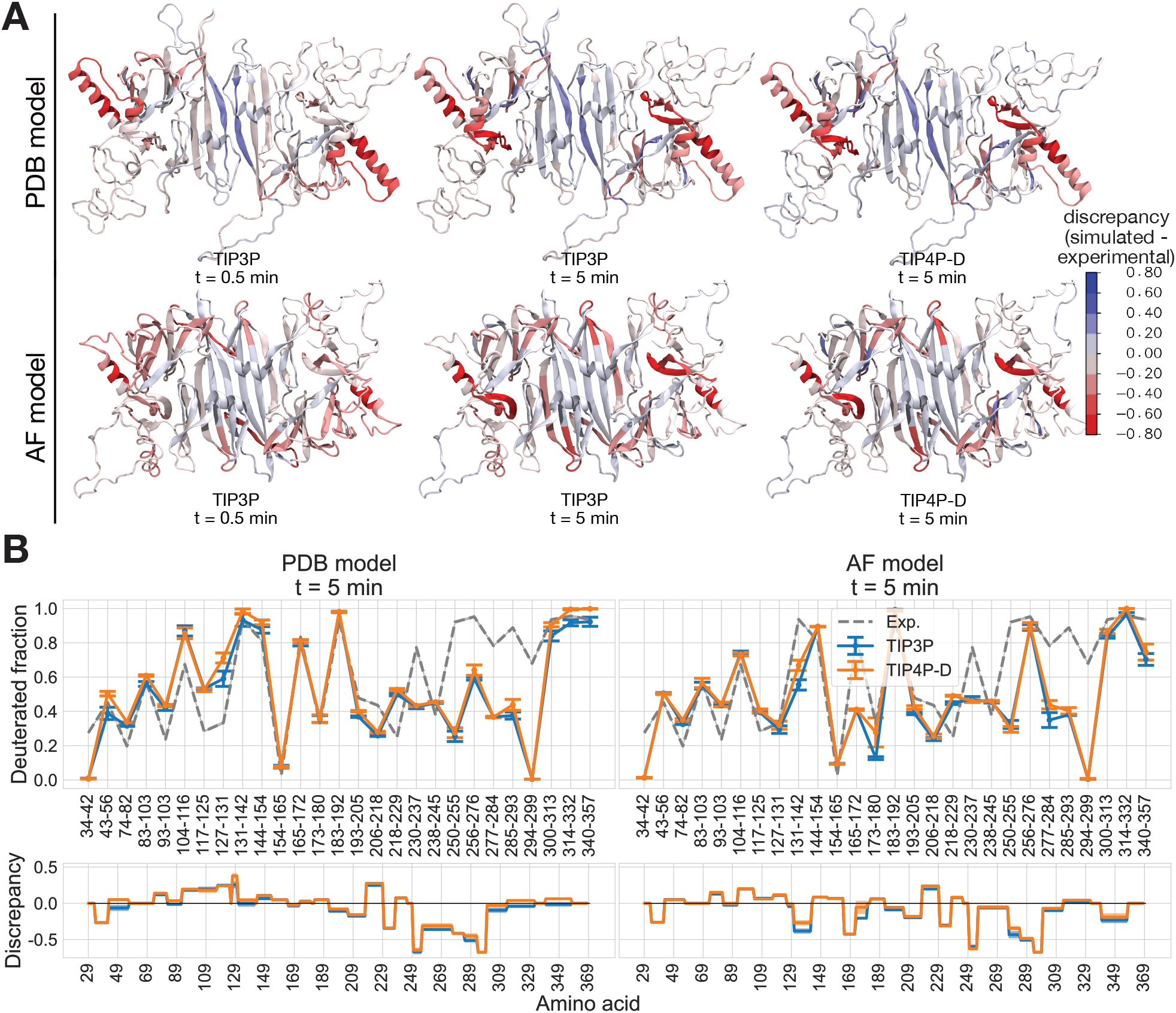
(A) Visualization of the discrepancy values from Fig. 1C for TIP3P and TIP4P-D water at incubation times of 0.5 min and 5 min, mapped onto the molecular structures of the PDB and AF models. Time points correspond to experimental incubation times, not MD simulation time. Shades of blue indicate regions where the simulated structure is more flexible and solvent-accessible than observed experimentally, while shades of red indicate regions where the simulated structure is more rigid and less solvent-accessible than expected. (B) Comparison of the deuterated fraction obtained from experimental results (dashed line) published by Amin-Wetzel et al.^25^ and the fraction computed from MD simulations (solid lines, blue for TIP3P water and orange for TIP4PD water, error bars indicate the standard deviation obtained from bootstrapping) for the PDB and AF model at incubation time point 5 min. Each point represents the mean value derived from three replicas and two monomers per replica. The error bars were obtained from bootstrapping. Below each absolute value plot, we report the discrepancy, which is defined as the difference between the simulated and experimental deuterated fractions.

**Figure 11.**
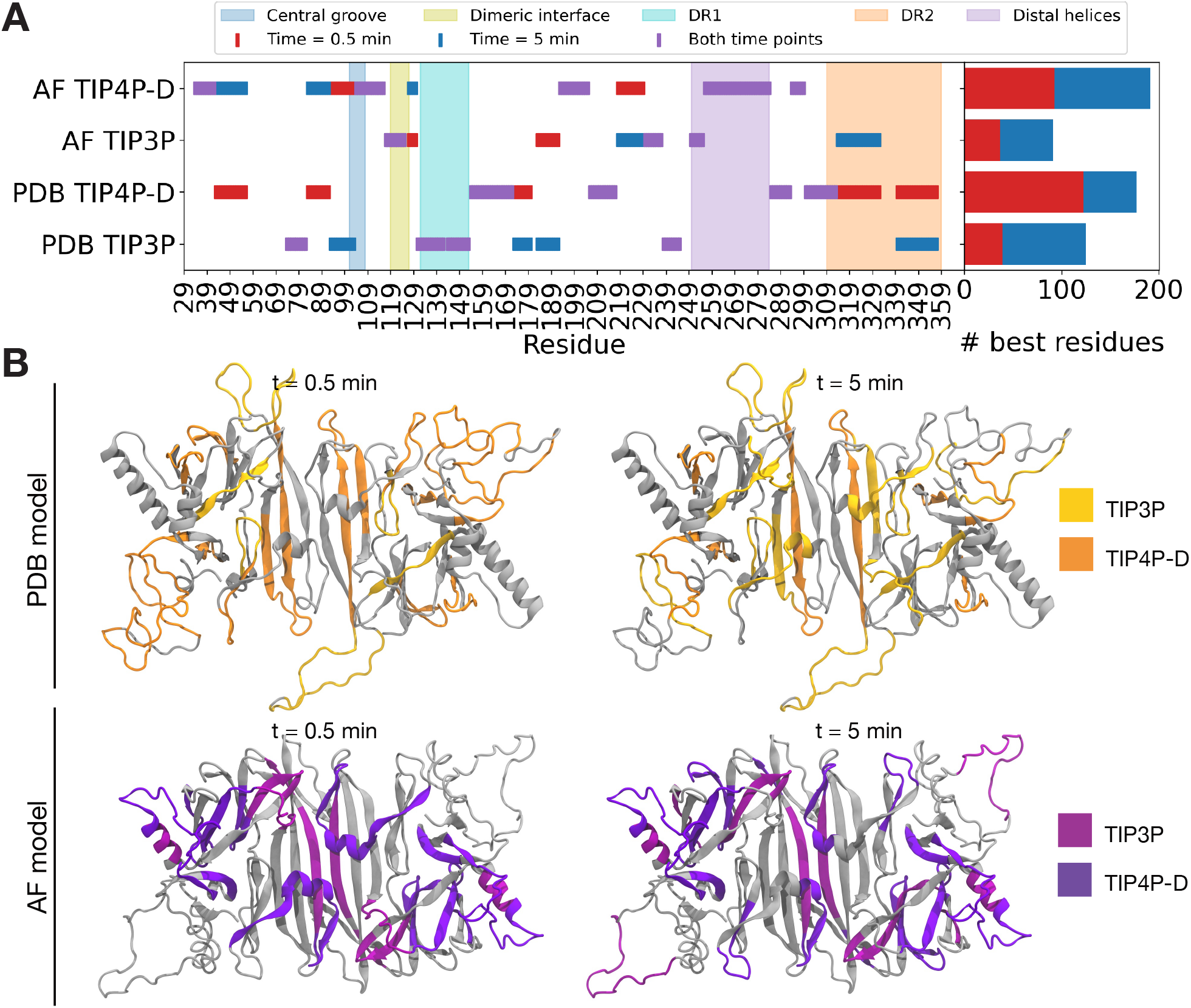
(A) Comparison of the deuterated fraction from Amin-Wetzel et al. ^25^ with the deuterated fraction computed from MD simulations (See Methods section ‘HDX data comparison’). For each residue, the plot identifies the model that best agrees with experimental data (‘best structure’) at each time examined point (t = 0.5 min, 5 min, or both). The histogram displays the cumulative number of residues for which each model most closely matches experimental data. Each of the four model variants is evaluated residue-wise at both time points, earning a point if it better reproduces the experimental deuterated fraction for a given residue compared to the other three models. The total score, obtained by summing points for each model at both time points, is represented in the histogram. (B) Best structural model of hIRE1*α* cLD dimer: colors indicate the best structural representation of each residue at either time point, as shown in Supplementary Fig.11A.

**Figure 12.**
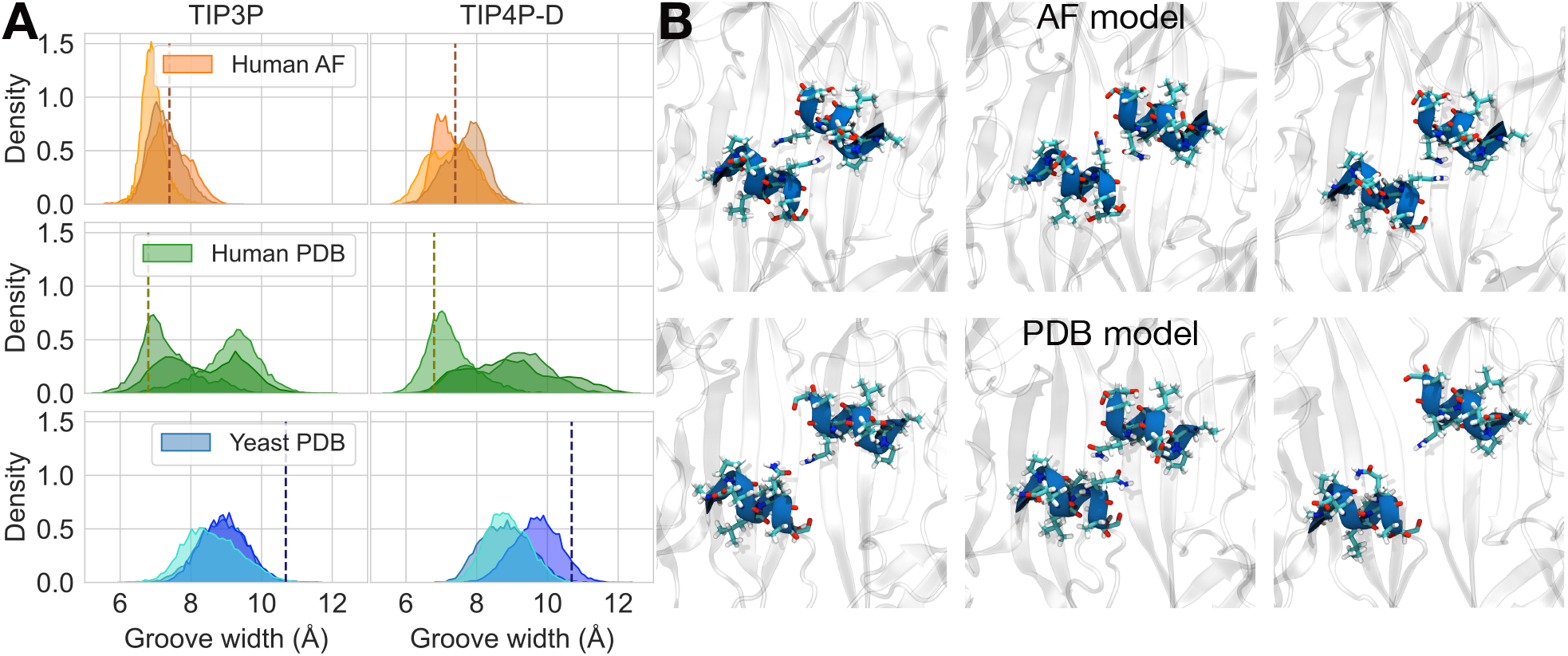
(A) Probability density distribution of the groove width of: hIRE1*α* cLD dimer (AlphaFold model) computed between C*α* of Q105 during simulations in TIP3P and TIP4P-D water (top panel); hIRE1*α* cLD dimer (PDB model) computed between C*α* of Q105 during simulations in TIP3P and TIP4P-D water (middle panel); yIre1 luminal domain dimer computed between C*α* of S200 during simulations in TIP3P and TIP4P-D water (bottom panel). Dashed lines indicate the groove width of the dimer model prior to energy minimization. (B) Representative Q105 side chains arrangements during simulations of the human cLD dimer AlphaFold model (top row) and of the PDB model (bottom row), snapshots are taken from three independent replicas in TIP3P and TIP4P-D.

**Figure 13.**
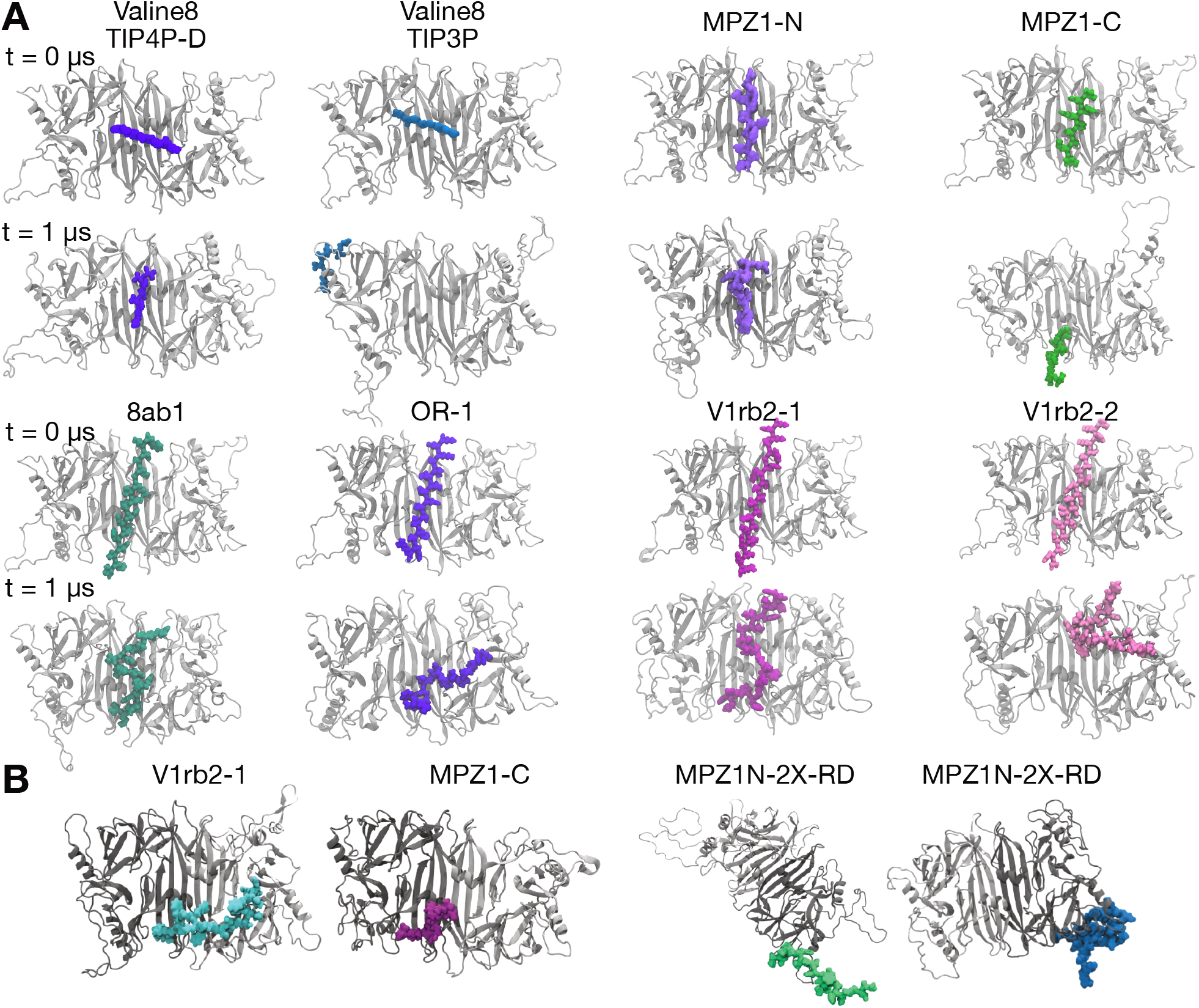
(A) Additional examples of peptide binding. Here, we report snapshots from simulations at the initial timestep (t= 0 μs) and final timestep (t= 1 μs) of the unfolded peptides not shown in Fig. 3A-C, in order: Valine8, MPZ1-N (perpendicular placement), MPZ1-C, 8ab1, OR-1, V1rb2-1, V1rb2-2. (B) Heterogeneous binding. Peptides that are unstable in the central groove placement tend to rearrange to different parts of IRE1, either to the side of the groove as for V1rb2-1 and MPZ1-C or to the outer part of the dimer in the proximity of the *α*B-helices, as in the case of MPZ1N-2X-RD.

**Figure 14.**
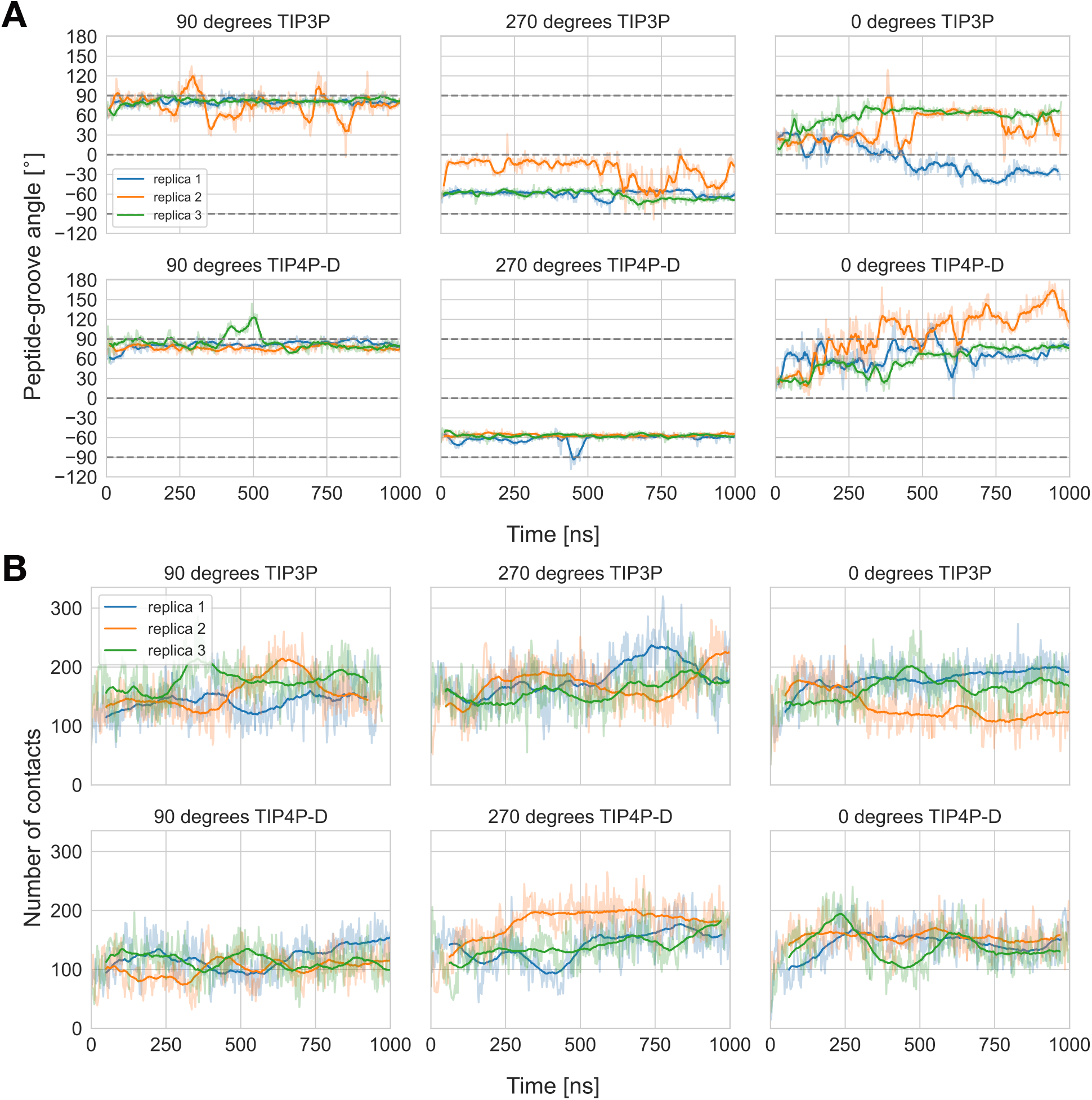
(A) Peptide orientation with respect to the central groove principal axis. The angle was computed as the dihedral angle described by the C*α* atoms of Y161 residues (groove principal axis) and the C*α* atoms of residues L1 and A12 of the MPZ1N peptide. The dark lines indicate the rolling average of the fraction of native contacts over 10 frames, while the shaded lines indicate the value per frame. (B) Number of contacts between hIRE1*α* cLD dimer and MPZ1N peptide. The dark lines indicate the rolling average of the fraction of native contacts over 50 frames, while the shaded lines indicate the value per frame. The analysis were performed on three sets of simulations: “90 degrees” orientation, the peptide is initially placed perpendicular to the central groove principal axis; “270 degrees” orientation, the peptide is initially placed perpendicular to the central groove principal axis but flipped 180 degrees with respect to the 0 degree; “0 degrees” orientation, the peptide is placed parallel to the groove principal axis.

**Figure 15.**
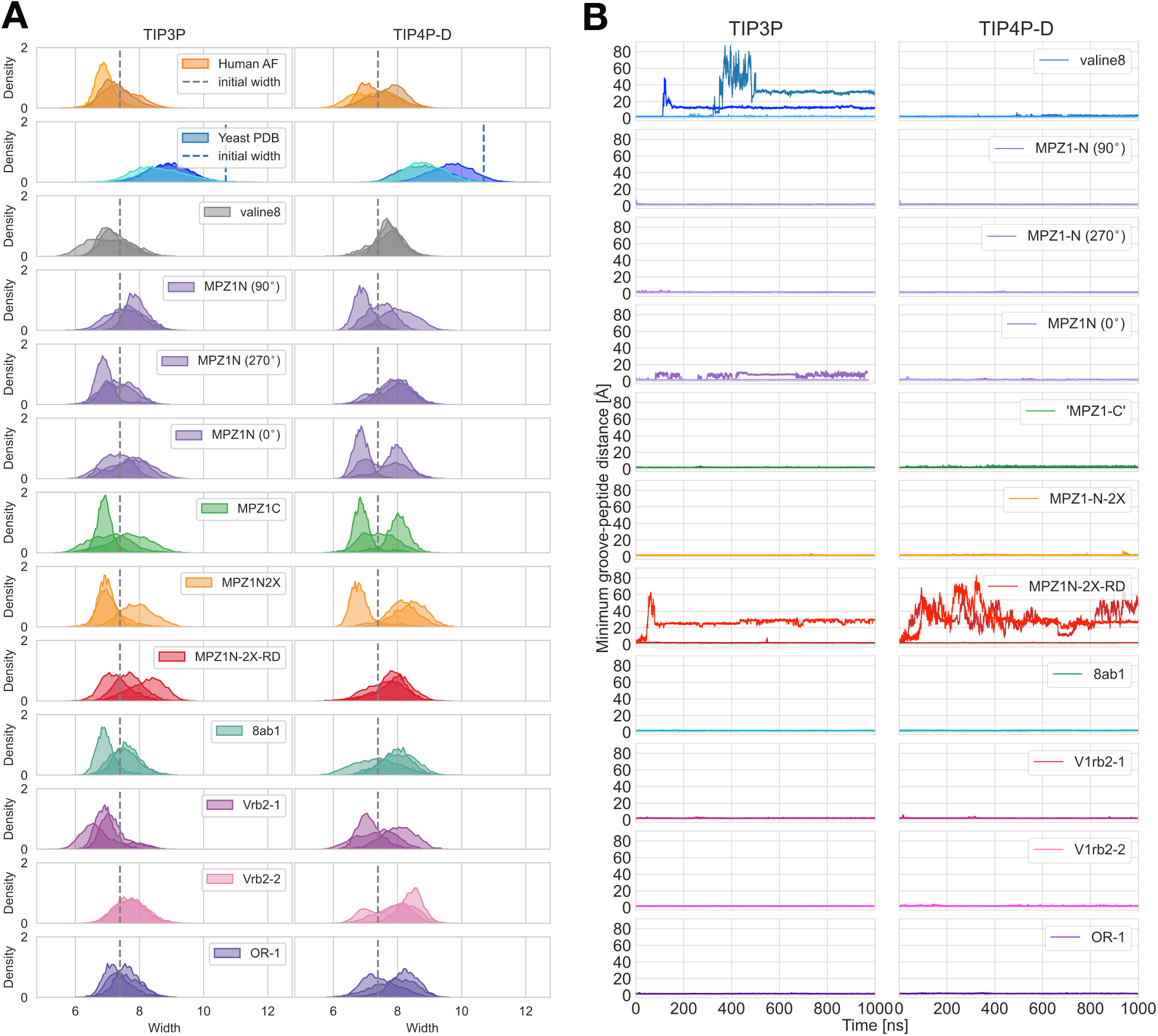
(A) Distributions of the groove width of peptide-bound cLD dimers throughout all simulations performed. The left column shows the values for the three replicas in TIP3P water, while the right column displays those for the three replicas in TIP4P-D water. (B) Minimum groove-peptide distance over time for all simulations of cLD dimer in complex with a peptide. The left column shows the values for the three replicas in TIP3P water, while the right column displays those for the three replicas in TIP4P-D water.

**Figure 16.**
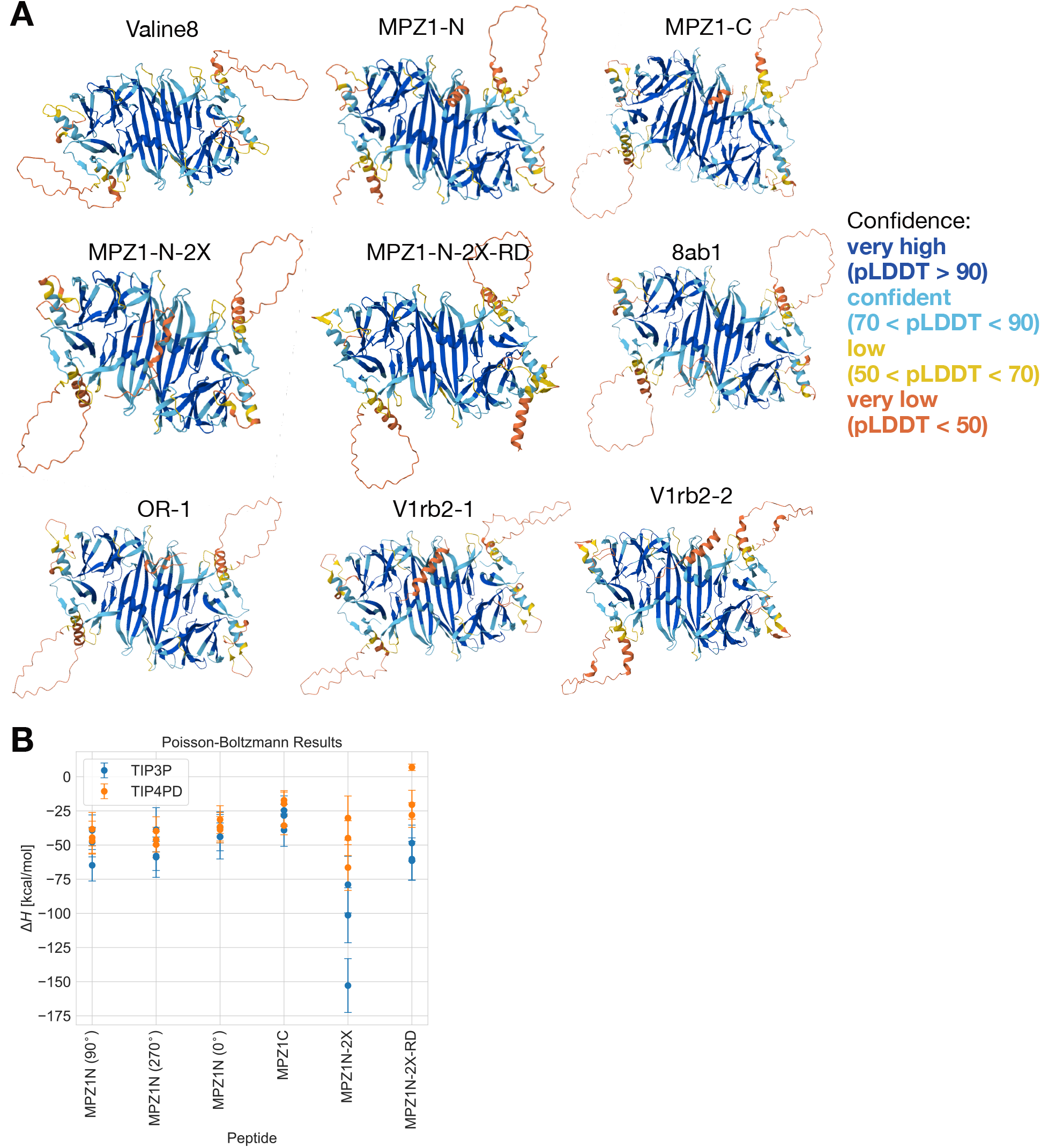
(A) Prediction of AlphaFold 3 for hIRE1*α* cLD dimer in complex with peptides. Colors represent the confidence of the prediction (plDDT). (B) Difference in enthalpy (enthalpy of binding, Δ*H*) as an estimate of the binding free energies of unfolded polypeptides to hIRE1*α* cLD dimer derived from MM/PBSA calculations of our peptide simulations.

**Figure 17.**
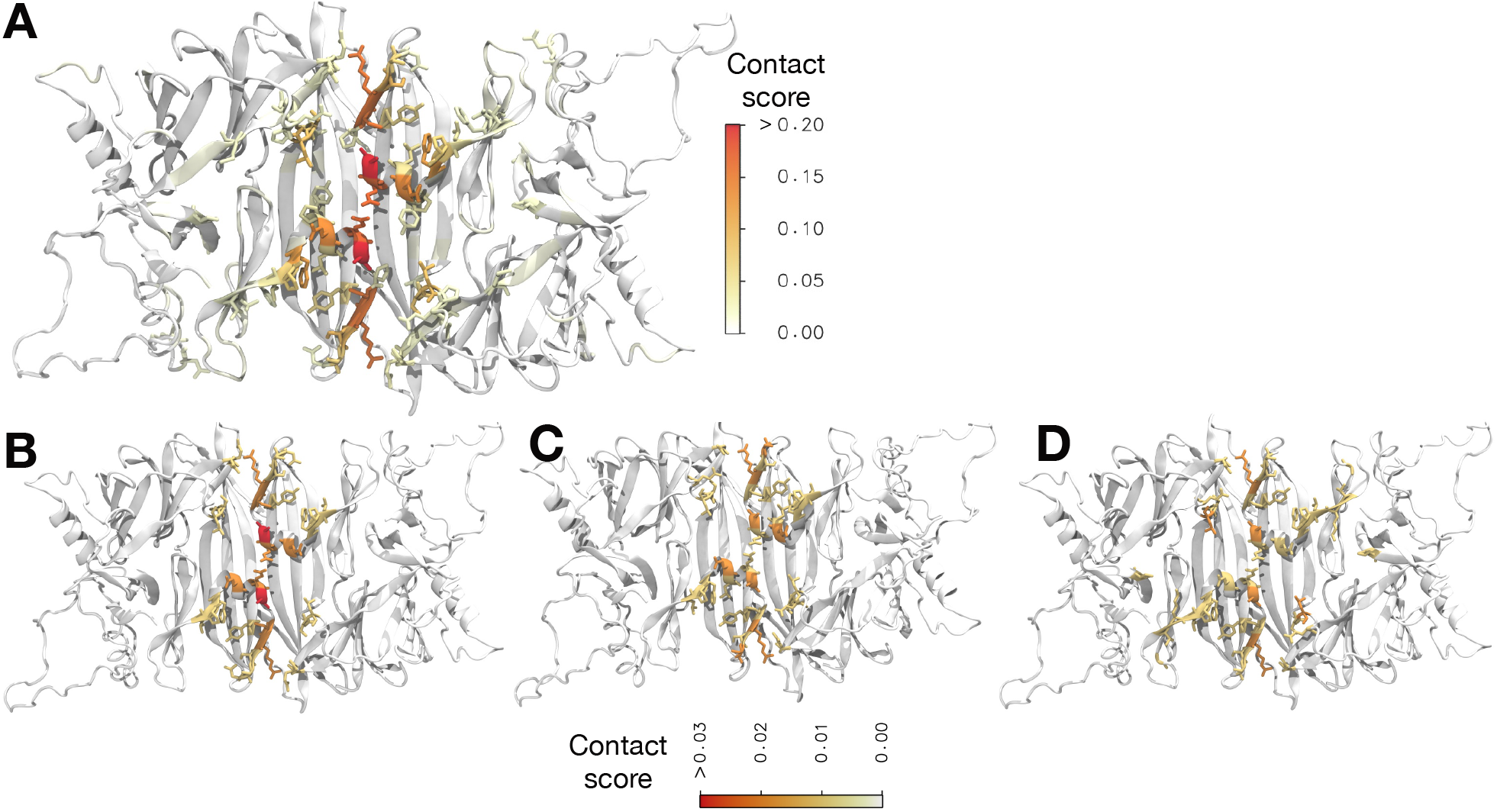
(A) Analysis of contacts between unfolded peptide and cLD dimer for all MD simulations performed in TIP3P water. The contact score reported on the AlphaFold model of the cLD dimer was obtained by computing the contacts from all the simulations, summing them, and normalizing them by the number of frames. Lower values indicate fewer contacts observed. (B) Contact analysis of MPZ1N-2X in complex with wild-type hIRE1*α* cLD dimer. (C) Contact analysis of MPZ1N-2X in complex with E102R hIRE1*α* cLD dimer. (D). Contact analysis of MPZ1N-2X in complex with Y161R hIRE1*α* cLD dimer. The analysis of panels B, C, and D took into consideration all six 1 μs-long replicas in TIP3P and TIP4P-D water for each system.

**Figure 18.**
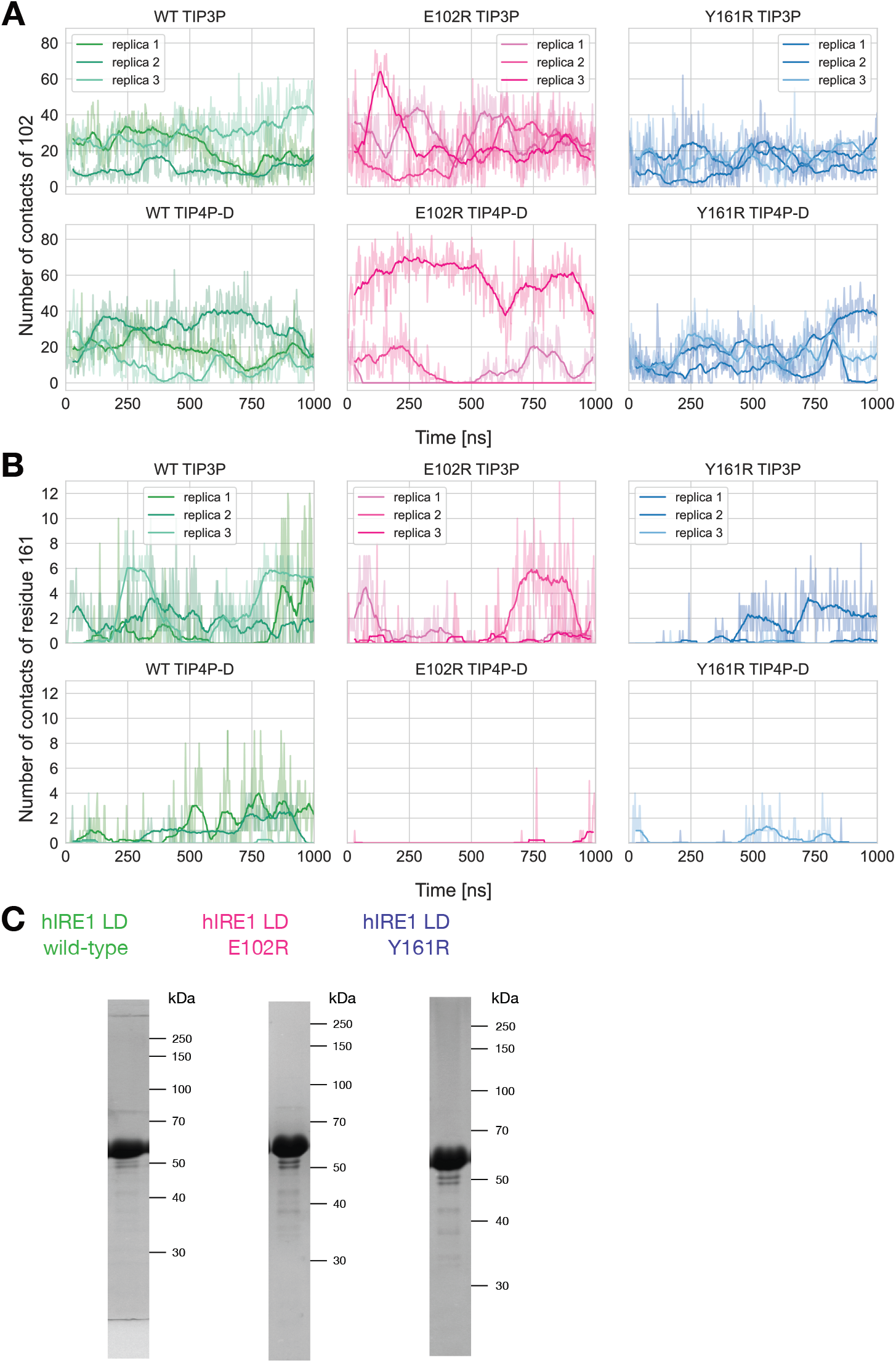
(A) Number of contacts between residues 102 on both monomers and the MPZ1-N-2X peptide during simulations of WT hIRE*α* LD and mutants E10R and Y161R. The dark lines indicate the rolling average of the fraction of native contacts over 25 frames, while the shaded lines indicate the value per frame. (B) Number of contacts between residues 161 on both monomers and the MPZ1-N-2X peptide during simulations of WT hIRE*α* LD and mutants E10R and Y161R. The dark lines indicate the rolling average of the fraction of native contacts over 25 frames, while the shaded lines indicate the value per frame. (C) Protein purification of WT hIRE*α* LD and mutants E10R and Y161R.

**Figure 19.**
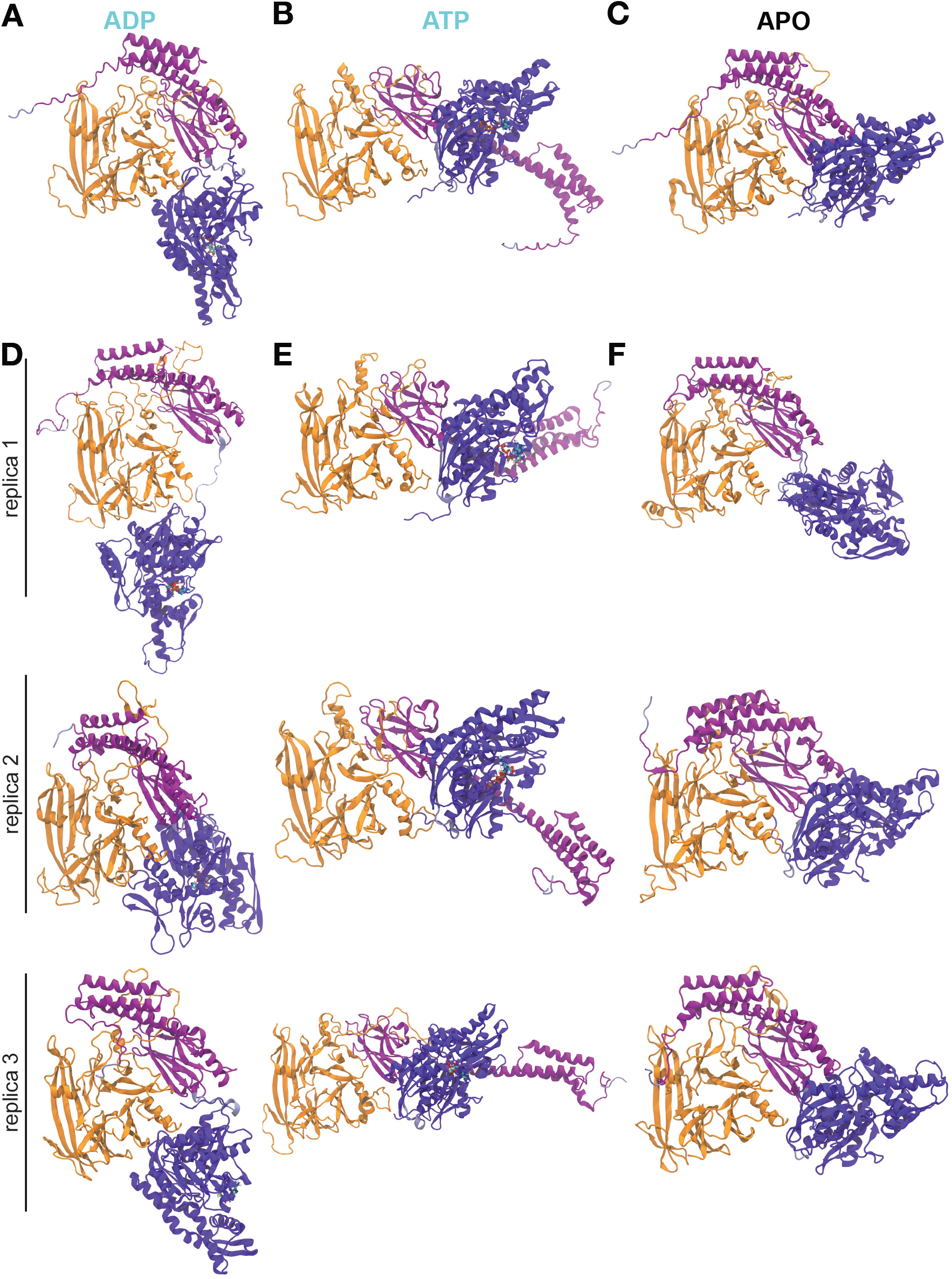
(A) Prediction of AlphaFold 3 for hIRE1*α* cLD monomer in complex with ATP-bound BiP. The colors are as in Fig. 5B. (B) Prediction of AlphaFold 3 for hIRE1*α* cLD monomer in complex with ADP-bound BiP. (C) Prediction of AlphaFold 3 for hIRE1*α* cLD monomer in complex with BiP not bound to any nucleotide. (D) Structure of hIRE1*α* cLD-BiP-ATP after 2 μs of simulation. (E) Structure of hIRE1*α* cLD-BiP-ADP after 2 μs of simulation. (F) Structure of hIRE1*α* cLD-BiP after 2 μs of simulation.

**Figure 20.**
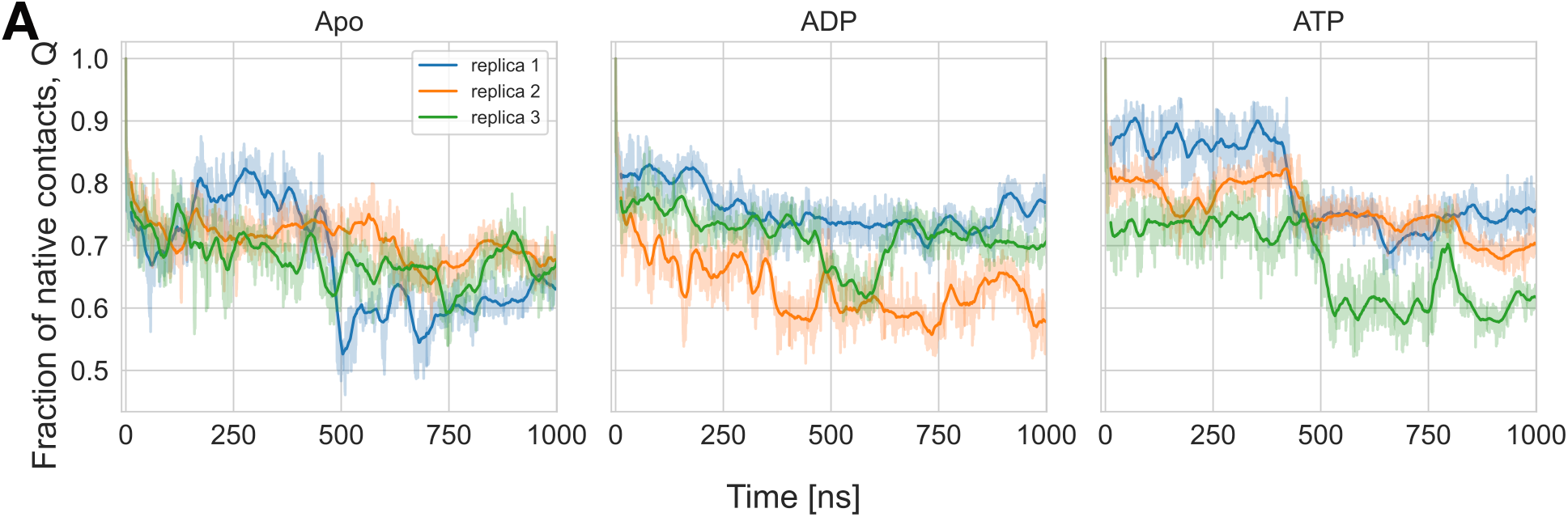
Fraction of native contacts between BiP and cLD monomer in simulations of the structures predicted by AlphaFold 3 without ligands or in complex with ADP or ATP. The dark lines indicate the rolling average of the fraction of native contacts over 100 frames, while the shaded lines indicate the value per frame. The fraction of native contacts (Q) was calculated according to the definition of Best et al.: ^67^ 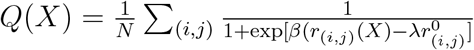 For *N* pairs of native contacts (*i, j*), where 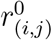 is the distance of the pair in the initial configuration (here the AlphaFold 3 prediction), *r*_(*i,j*)_(*X*) is the distance at frame X, *β* is a smoothing parameter (*β* = 50 *nm*^−1^), *λ* is the tolerance of the reference distance (*λ* = 1.8) and the cutoff used to define a contact between heavy atoms was 0.45 nm.

